# Global Evaluation of Congenital Heart Disease-Associated Non-Coding Variants

**DOI:** 10.64898/2025.12.02.691900

**Authors:** Edwin G. Peña-Martínez, Shreya Sharma, Joshua G. Medina-Feliciano, Elise Root, Lois G. Parks, Marissa Granitto, Diego A. Pomales-Matos, Jean L. Messon-Bird, Adriana C. Barreiro-Rosario, Leandro Sanabria-Alberto, Alejandro Rivera-Madera, Jessica M. Rodríguez-Ríos, Rosalba Velázquez-Roig, Juan A. Figueroa-Rosado, Mackenzie Noon, Omer A. Donmez, Carmy Forney, Hayley K. Hesse, Katelyn A. Dunn, Xiaoting Chen, Matthew R. Hass, Lucinda P. Lawson, Matthew T. Weirauch, Leah C. Kottyan, Steven K. Reilly, Devesh Bhimsaria, José A. Rodríguez-Martínez

## Abstract

Genome-wide association studies (GWAS) have mapped thousands of congenital heart disease (CHD)-associated variants within non-coding regions of the genome. Non-coding variants can alter regulatory mechanisms, such as transcription factor (TF) binding control of gene expression, potentially contributing human diseases. However, with the increasing number of disease-associated variants, comprehensive functional validation remains a significant challenge. In this work, we developed a novel method called SNP Bind-n-Seq to evaluate >3,000 CHD-risk variants for allelic binding for the cardiac TFs NKX2-5, GATA4, and TBX5 in a high-throughput manner. These binding affinity data sets were coupled with a massively parallel reporter assay (MPRA) to screen CHD-risk variant genotype-dependent regulatory activity. We identified 170 variants that exhibit allelic TF binding and 187 that modulate gene expression. Combining both approaches revealed three high-confidence variants with genotype-dependent TF binding, genotype-dependent transcriptional activity, and eQTL behavior in cardiac cells. Collectively, this study provides the first combined high-throughput biochemical and functional genomic evaluation of thousands of CHD-risk variants.

**Highlights:**

- Allelic binding affinity measurements of ∼9,600 variants for NKX2-5, GATA4, and TBX5
- EvaluaFon of >3,000 CHD-risk variants for genotype-dependent regulatory acFvity
- InteracFon networks idenFfy funcFonal variants and genes involving cardiac eQTLs

**Graphical Abstract:** 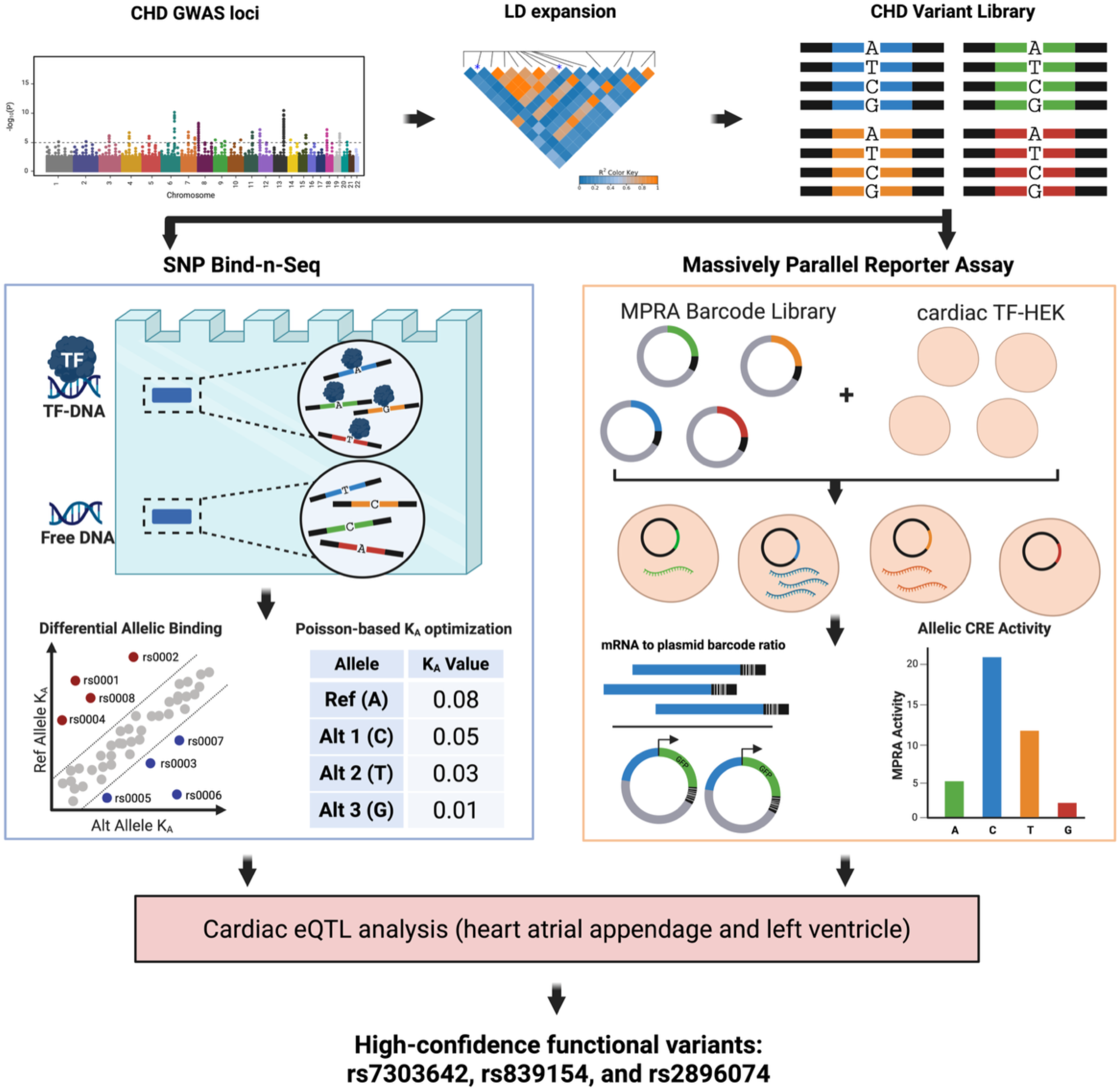

## Introduction

Congenital heart diseases (CHDs) are the most common birth defect, affecting 1 in every 100 liveborn infants. CHDs are characterized by structural abnormalities in the heart and vessels that cause thousands of deaths each year, mainly in infants under one year of age. ^1–3^ Among the multiple causes of CHDs are genetic variants that alter regulatory mechanisms during heart development. ^4–6^ Genome-wide association studies (GWAS) have identified hundreds of CHD-associated variants, with the overwhelming majority (>90%) mapping to non-coding regions of the genome. ^7–11^ Non-coding genetic variants, such as single nucleotide polymorphisms (SNPs), have been suggested to contribute to CHD etiology by altering the functions of cardiac transcription factors (TFs), which bind to DNA to regulate cardiac differentiation. ^2,12–16^

NKX2-5, GATA4, and TBX5 are evolutionarily conserved cardiac TFs necessary for heart development in vertebrates, and mutations impairing their function have been implicated in multiple types of CHDs. ^12,17–23^ For example, previous work has identified coding variants within their DNA-binding domains (DBDs) to be causal for CHDs. ^21,24–30^ Each of these cardiac TFs belongs to its own TF family: Homeobox, GATA, and T-box family, respectively, with each TF displaying unique DNA binding preferences. ^18,31^ NKX2-5, GATA4, and TBX5 work synergistically and cooperatively to regulate cardiac genes needed to initiate cardiogenesis and differentiation of multiple tissues within the heart. ^24,28,32–34^ Non-coding variants alter the DNA-binding affinity of cardiac TFs, potentially leading to multiple types of cardiovascular diseases. ^8,15,20,35–44^ However, with a continuously increasing number of CHD-risk variants being discovered, testing each of them to identify functional variants remains challenging.

In this work, we systematically evaluated over 3,000 CHD-risk variants for allele-biased regulatory function involving cardiac TF-DNA binding and transcriptional activity. Building upon previous methods ^45,46^, we developed SNP Bind-n-Seq, a high-throughput gel shift-based assay to quantify differential allelic binding events. Using this assay, we constructed allelic enrichment curves and, through a Poisson-based optimization framework, derived binding affinities for the cardiac TFs NKX2-5, GATA4, and TBX5. To assess gene regulatory effects, we also performed a massively parallel reporter assay (MPRA) ^47–50^ to identify CHD-risk variants that exhibited genotype-dependent transcriptional activity. Using a combined experimental and computational approach, we identified 352 CHD-risk variants with allelic regulatory functions for either cardiac TF binding (170 variants) or *cis*-regulatory element (CRE) activity (187 variants). Additionally, we identified three high-confidence variants with an allelic skew in both TF binding and transcriptional activity, and are in cardiac expression quantitative trait loci (eQTL), which can be further explored for roles in CHD disease mechanisms. In summary, this study provides the first combined biochemical and functional genomic evaluation of thousands of CHD-risk variants, offering a scalable approach that can be applied broadly to complex human diseases.

## Results

### Quantifying allelic cardiac TF binding through SNP Bind-n-Seq

To evaluate the impact of non-coding variants on cardiac TF binding, we first collected all CHD-associated variants from the GWAS catalog ^7^ available in November 2022. We identified 121 CHD-associated SNPs from six GWASs ^51–56^ which were expanded to include variants in linkage disequilibrium (LD) in four ancestries (EUR, AFR, EAS, SAS; R^2^ > 0.80), totaling 3,232 unique variants. An oligonucleotide library of the 3,232 variants containing every possible variant allele was synthesized (12,928 unique sequences, **Supplementary Data 1**). Variants were centered on a 40 bp DNA sequence of genomic context with constant regions for downstream barcoding and sequencing (102 bp total, **Supplementary Figure 1A**).

To evaluate the impact of CHD-risk variants on TF-DNA binding, we developed SNP Bind-n-Seq, a high-throughput gel shift-based assay coupled to DNA sequencing (**Figure 1A**). In SNP Bind-n-Seq, a TF is equilibrated with the oligonucleotide library containing thousands of sequence variants. TF-bound and -unbound sequences are separated in a native polyacrylamide gel, and both fractions are sequenced. SNP Bind-n-Seq was performed with the purified DBD of NKX2-5, GATA4, and TBX5 at seven concentration points ranging from 0 nM to 3,000 nM. Experiments were performed in duplicates for each TF. The oligonucleotide library was modified to have a fluorescent probe through primer extension reaction as previously described ^57^, which allowed gel excision of bound and unbound fractions at each concentration. Bound and unbound fractions were individually barcoded with a unique identifier for pooling and downstream computational analysis.

**Figure 1:**
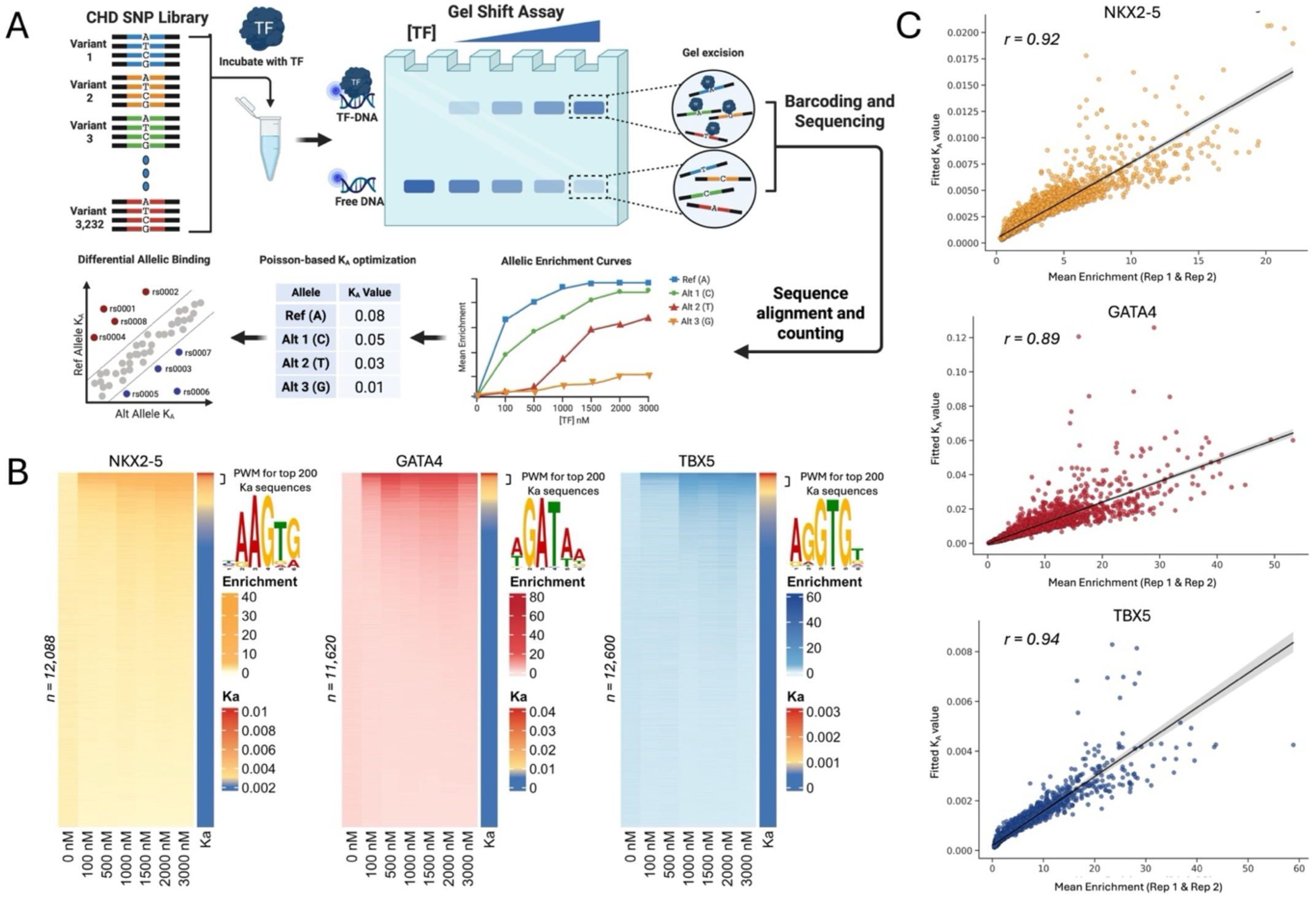
High-throughput evaluation of TF binding through SNP Bind-n-Seq. **A)** Overview of SNP Bind-n-Seq experimental approach and computational analysis. **B)** Sequence enrichment and binding affinity measurements for CHD-associated variants. Enrichment was calculated for all sequences at seven concentration points ranging from 0 nM to 3,000 nM for NKX2-5 (left), GATA4 (middle), and TBX5 (right). PWM logos generated from the top 200 sequences with the highest binding affinity (KA). **C)** Correlation between mean enrichment scores and fitted KA values. Enrichment at 3,000 nM is displayed for NKX2-5 (top), GATA4 (middle), and TBX5 (bottom).

To quantify TF binding through SNP Bind-n-Seq, we employed a statistical framework for all CHD-risk variants centered within the 40-bp sequence. Assuming non-cooperative, site-independent binding, we compute occupancy probabilities based on relative affinities at varying protein concentrations. To estimate these affinities, a joint model fits bound and unbound read counts across concentrations, inferring a concentration-independent binding parameter KA for each sequence (see Method for details). KA is conceptually analogous to a binding constant, which is defined as the molecular interaction strength at equilibrium. Read counts were modelled as a Poisson distribution, and KA is optimized to maximize the likelihood of the observed data. Enrichment was then calculated by comparing bound counts to a pseudo-unbound estimate, derived by scaling total unbound reads in each replicate by a factor of 2.5 and normalizing to a million. The resulting KA values were used to reconstruct expected counts and enrichment across concentrations.

Using this approach, we quantified TF binding enrichment and affinity for all 12,928 sequences (3,232 base-permutated variants; **Figure 1B**). Duplicates across all concentration points, excluding 0 nM, showed strong correlation for all three TFs (R^2^ > 0.75), indicating strong experimental reproducibility (**Supplementary Figure 1B and C**). TF motif derivation for the top 200 bound sequences produced position weight matrix (PWM) logos similar to those previously described for NKX2-5, GATA4, and TBX5 (**Figure 1B, Supplementary Figure 2**). ^15,16^ Additionally, the Ka values showed strong correlation with the TF binding enrichment measurements for NKX2-5 (r = 0.92), GATA4 (r = 0.89), and TBX5 (r = 0.94; **Figure 1C**). Collectively, these results indicate that SNP Bind-N-Seq experimental data are highly reproducible and recapitulate known TF DNA binding specificities.

### SNP Bind-n-Seq identifies CHD-risk variants with allele-biased TF binding

Using SNP Bind-n-Seq, we constructed allelic enrichment curves and quantified binding affinity for 3,232 CHD-risk variants for all three cardiac TFs. In doing so, we identified 170 CHD-risk variants (∼5% of all tested variants) with >2-fold differential allelic binding compared to the reference genome allele for NKX2-5 (54 SNPs; 31 increase, 23 decrease), GATA4 (58 SNPs; 30 increase, 28 decrease), or TBX5 (62 SNPs; 35 increase, 27 decrease) (Figure 2A and **Supplementary Figure 3A**). As controls, we included variants with differential binding for NKX2-5 that we previously described through electrophoretic mobility shift assay (EMSA), and observed a significant correlation (R^2^ = 0.95, *p*-value = 0.012, **Supplementary Figure 3B**). ^35^

**Figure 2:**
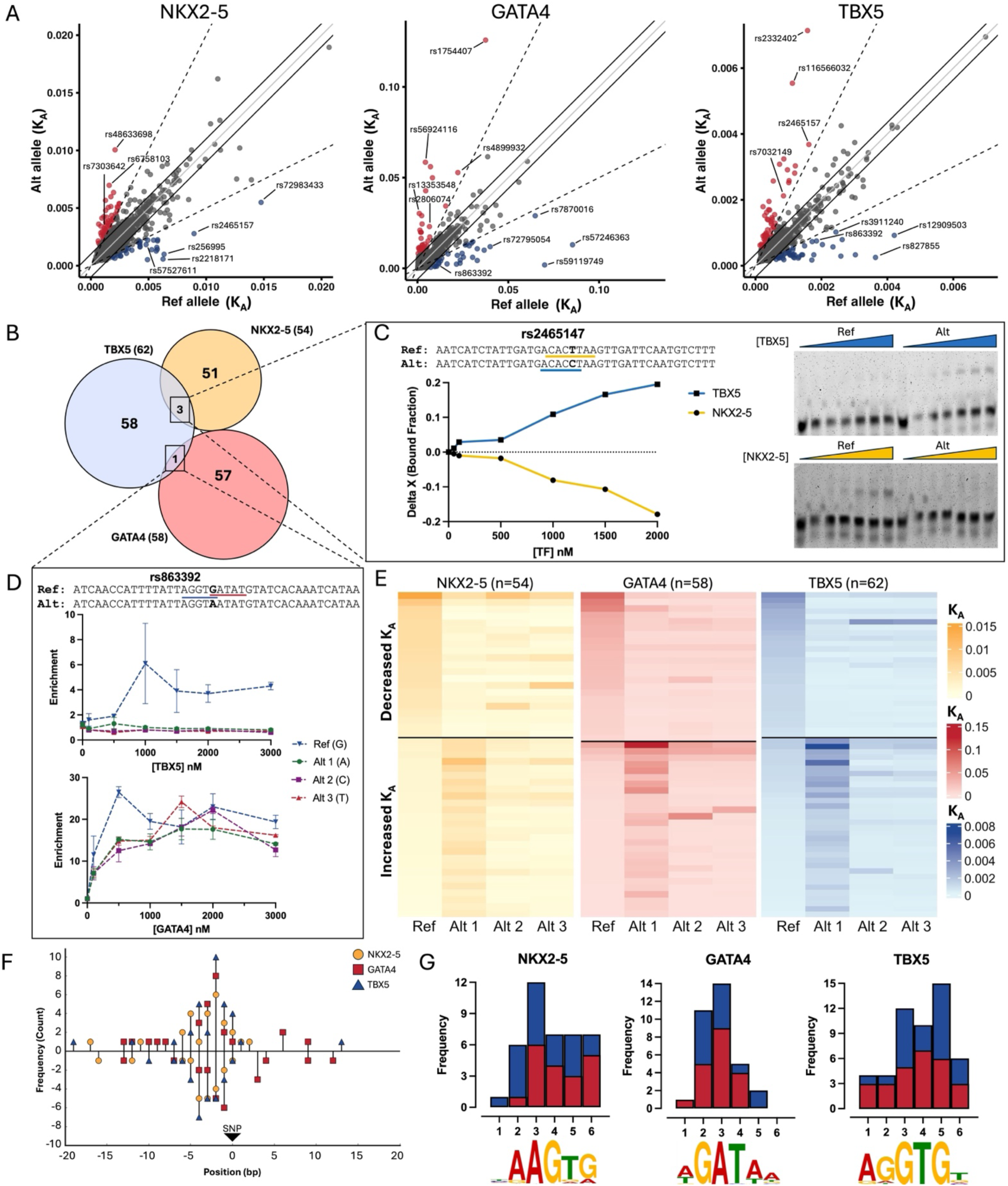
CHD-associated variants exhibit allele-biased binding for cardiac TFs. **A)** Differential binding affinity analysis between reference and CHD-risk alleles. Variants with increased binding affinity (higher KA value for the alternate allele) are represented in red, while a decrease in binding affinity (higher KA value for the reference allele) is represented in blue. The solid gray line represents the Y = X intercept with a slope of 1. The dashed line has a 15° angle from the solid line and represents a 2-fold change in binding affinity between the reference and alternate allele. **B)** Venn diagram of CHD-risk variants with differential allelic binding. Overlaps between diagrams represent variants that altered DNA-binding for multiple TFs. **C)** *In vitro* validation of rs2465147 through EMSA for TBX5 (top) and NKX2-5 (bottom). Binding sites in the reference sequence are underlined in yellow for NKX2-5 and in blue for TBX5 in the alternate sequence. **D)** Allelic enrichment curve of rs863392 for TBX5 (top) and GATA4 (bottom). Reference alleles (Ref) are represented in blue, and tag-SNP alleles from the GWAS catalog (Alt 1) are represented in green. Permutated alleles (alternate non-risk; Alt 2 and Alt 3) are represented in red and purple, respectively. **E)** Heatmaps illustrating allele-specific PrOBEX fitted K-values for SNPs predicted to alter transcription factor binding of NKX2-5 (left), GATA4 (center), and TBX5 (right). Each row corresponds to an individual SNP (rsID), with columns representing the reference allele and all possible alternative nucleotides. “Alt1” denotes the observed alternative allele reported in the GWAS catalog, while “Alt2” and “Alt3” correspond to the remaining permutated alleles. Cell color intensity reflects the magnitude of the fitted K-value, with warmer colors indicating stronger predicted binding affinity. The upper and lower panels display SNPs associated with decreased and increased binding affinity, respectively. **F)** Distribution of TF binding motifs relative to the position of the SNP with allelic binding. Dots represent the number of motifs created or disrupted for NXK2-5 (yellow circles), GATA4 (red square), and TBX5 (blue triangle). The X-axis represents genomic coordinates, a 40 bp window in the SNP-Bind-n-Seq assay. The arrow represents the SNP location at X = 0. **G)** Nucleotide contribution of variants that directly create or disrupt TF binding motifs for NKX2-5 (left), GATA4 (middle), and TBX5 (right). The contribution of created motif counts is presented in red, and disrupted motifs are shown in blue. The motif used to scan variant contribution is displayed below the X-axis, where the value represents position within the motif. The bars in the plot are overlapping, not stacked.

Next, we proceeded to evaluate biochemical mechanisms that could be driving allelic binding events of cardiac TFs. We identified four variants (rs863392, rs2465147, rs28394479, and rs77931854) that altered the binding of two TFs (**Figure 2B**). For example, variant rs2465147(T>C) had a significant decrease in NKX2-5 binding affinity, but an increase for TBX5. When observing the risk (C) and non-risk (T) alleles, variant rs2465147 disrupts the NKX2-5 binding motif (5’-CAC<u>T</u>T −3’) while simultaneously creating a consensus TBX5 binding motif (5’-ACAC<u>C</u>T −3’) (**Figure 2C, Supplementary Figure 4**). Likewise, variant rs863392(G>A) disrupted two binding motifs, decreasing the binding affinity of TBX5 (5’-AGGT<u>G</u>T −3’) and GATA4 (5’-A<u>G</u>ATAA −3’) (**Figure 2D**). Our experiment included all four nucleotides in the central base, providing additional information on alleles with no disease or trait association to date. We observed multiple variants where an alternate non-risk allele had a larger effect on TF-DNA binding (e.g., rs57527611 for NKX2-5, rs13353548 for GATA4, and rs3911240 for TBX5, **Figure 2E; Supplementary Figure 5**).

Finally, we explored biochemical mechanisms behind SNPs with allelic binding. Specifically, we asked whether variants with allelic binding directly create or disrupt TF binding motifs. From the 170 SNPs that exhibited allelic differences in binding in the SNP Bind-N-Seq experiments, 126 (74%) directly created (71/126, 56%) or disrupted (55/126, 44%) binding motifs for one of three tested TFs (**Figure 2F, Supplementary Figure 6**). The remaining 44 variants (26%) did not create or disrupt a core TF binding motif. Some variants with differential binding occurred in the flanking regions of the TF binding site, which have been previously shown to contribute to TF regulation. ^58,59^ Another subset of variants created low-affinity binding sites adjacent to core binding motifs of NKX2-5, GATA4, and TBX5. This suggests that for approximately one-quarter of variants that experimentally exhibit allelic binding, changes in TF binding are likely not predicted using motif-dependent models. However, for all three TFs, binding motifs were more likely to be created rather than disrupted (**Figure 2G**).

The experimental scope of this work was limited to only three TFs, which may not fully describe the complete regulatory mechanisms behind CHD etiology. To identify other TFs potentially impacted by the identified CHD-associated variants, we performed a HOMER motif discovery analysis to identify allelic TF binding sites that are being created or disrupted. From the 170 variants that exhibited allelic binding, we identified 931 unique TF binding motifs that occur in either the reference or risk allele (**Supplementary Figure 7A-B**). Of those, 602 (65%) created new TF binding motifs that were absent in the reference allele; the remaining 329 (35%) disrupted motifs in the reference allele and are absent in the alternate sequence. Motifs were created for 21 TF families (213 TF motifs) and disrupted for 18 TF families (133 TF motifs; **Supplementary Figure 7C**). These findings suggest that for the 170 SNPs identified in our work, motifs are more likely to be created than disrupted, which is consistent with our experimental results for NKX2-5, GATA4, and TBX5. As expected, when SNPs altered binding for NKX2-5, GATA4, and TBX5, we observed created and disrupted motifs from TFs of the same family (e.g., NKX2-2, NKX3-1, GATA3, GATA6, TBX6, TBX21, etc.), which are also involved in the GRNs of heart development (**Supplementary Figure 7D**). ^5,14–16,60^

### Computational prediction of GWAS SNPs on cardiac TF binding

To further evaluate the potential impact of disease-associated SNPs, we trained three computational models (MinSeqS ^61^, LS-GKM-SVM^62,63^, and MEME^64^) with the NKX2-5, GATA4, and TBX5 *in vitro* data generated from SNP Bind-n-Seq, and scored every variant from the GWAS catalog. ^7^ For this, we used the top 500 sequences with the highest KA values of each TF and trained the three models to predict changes in NKX2-5, GATA4, and TBX5 DNA binding (**Figure 3A**). ^61,62,64^ As a negative set we used random genomic sequences within the same chromosomes that match length and GC content. Performance parameters were determined by a 60-40 split of the dataset, training the model with 60% of the data and scoring the remaining 40% in multiple iteration to determine the area under the receiver operating curve (AUROC; Figure 3B and **Supplementary Figure 8A**).

**Figure 3:**
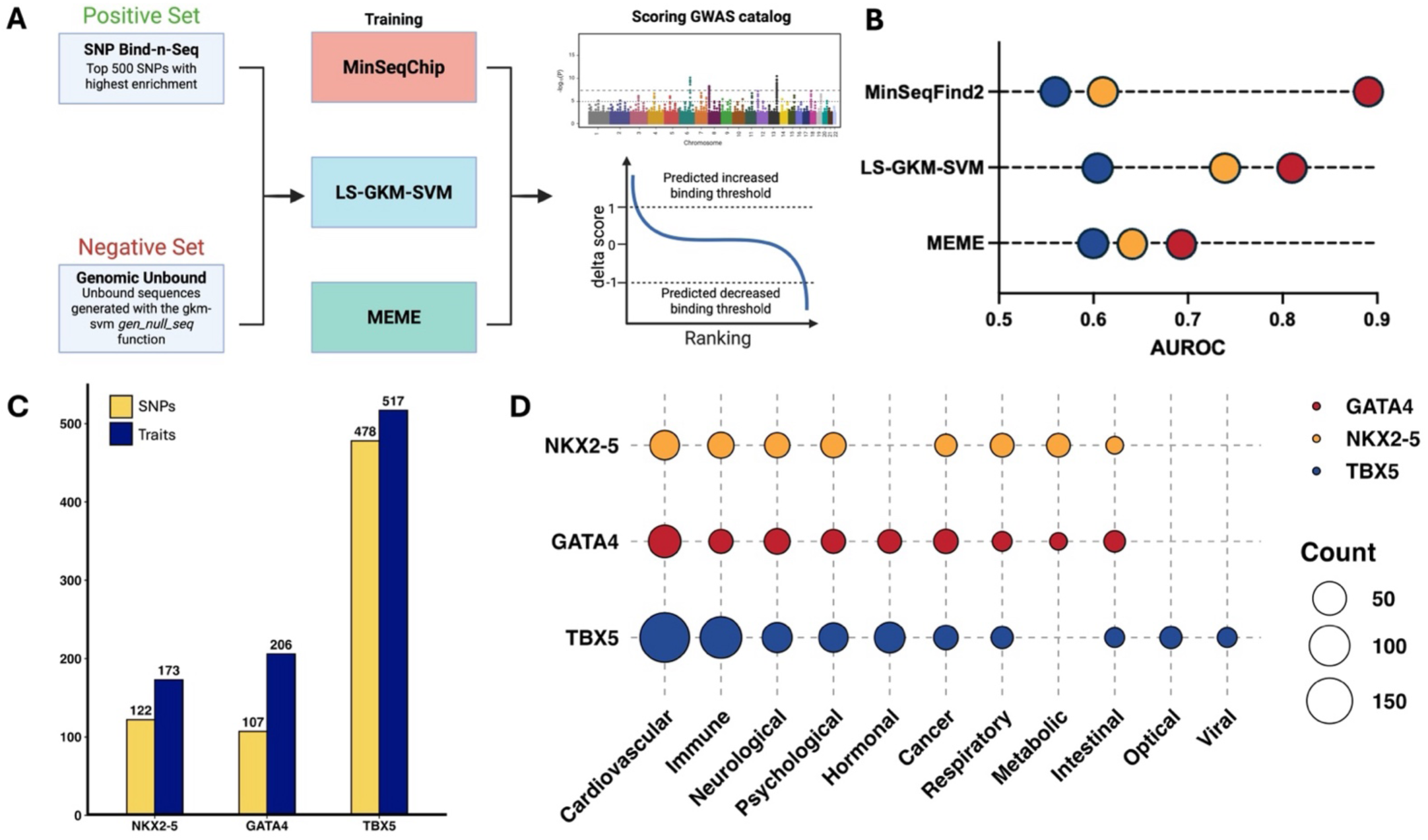
Computational prediction of GWAS catalog variants on cardiac TF-DNA binding. **A)** Schematic of model training using SNP Bind-n-Seq binding data. **B)** Performance parameters of three predictive models trained on SNP Bind-n-Seq binding data. **C)** Number of SNPs (yellow) with traits (blue) from the GWAS catalog predicted to alter NKX2-5, GATA4, and TBX5 binding. **D)** Number of disease-associated SNPs divided by trait parent term predicted to alter NKX2-5, GATA4, and TBX5 binding. Dots in the figure represent NKX2-5 (yellow), GATA4 (red), and TBX5 (blue).

After scoring all disease-associated SNPs from the GWAS Catalog (n = 235,773), we identified 709 variants predicted to alter NKX2-5, GATA4, or TBX5 DNA binding (**Figure 3C**). From those predicted SNPs, 243 (∼34 %, Odds ratio = 4.69, p-value = 3.3 x 10^-68^) were associated with cardiovascular diseases (CVDs) and traits, mainly blood pressure, CHDs, heart function/structure, and cardiac cell traits (**Figure 3D and Supplementary Figure 8B**). Variants with cardiovascular phenotypes had the highest counts for all three TFs, which is unsurprising considering their extensive research and contributions to cardiovascular trait mechanisms. ^17,21,22,30,36,65,66^ Additionally, we identified non-CVD associated variants, such as neurological and immune diseases and traits (**Supplementary Figure 8B**). Although these TFs are mainly studied in the context of cardiovascular genetics, there is evidence of their role in the regulatory mechanisms of other tissues. For example, GATA4 has been identified to play a key role in liver development and immune responses, and TBX5 has been shown to be involved in limb development. ^67–71^ These findings could open potential research areas towards establishing functional mechanisms of NXK2-5, GATA4, and TBX5 outside of CVDs and other previously identified developmental pathways.

### CHD-risk variants in cardiac regulatory elements have transcriptional activity

To identify CHD-risk variants regulating gene expression, we performed a massively parallel reporter assay (MPRA; **Supplementary Figure 9**) in Flp-In cell lines that were modified to stably express NKX2-5, GATA4, and TBX5 (**Supplementary Figure 10**). For the MPRA library, we compiled all CHD-risk associations (n = 157) listed in the GWAS catalog as of February 2024. For each of the tag SNPs, we performed an LD expansion (r^2^ ≥ 0.8) in each ancestry of the initial genetic association(s). In total, we identified 5,431 variants, totaling 14,524 unique sequences with every reported variant allele. Oligos of 170 bp centered on the variant were cloned, synthesized, and labeled with degenerate 20-mer barcode sequences using PCR followed by cloning upstream of an *eGFP* gene under the control of a minimal promoter.

For downstream analysis after performing MPRA, we considered variants with at least 10 unique barcodes, resulting in 14,114 oligos (97.2% of assayed variants; **Supplementary Figure 11A**). The normalized *eGFP* mRNA to plasmid control barcode ratio was used to quantify CRE activity driven by each oligonucleotide.

Using the finalized CHD MPRA library, we first identified variants capable of driving CRE activity. A variant was considered to have CRE activity if any of its alleles had a significant increase in transcriptional activity (*eGFP* mRNA: plasmid DNA ratio, **Supplementary Figure 11B**). Experiments were performed in triplicate with strong correlation of MPRA activity (log2 fold change RNA/DNA) between replicates (R^2^ >0.99, **Supplementary Figure 11C**). We therefore defined that a sequence in the MPRA library had CRE activity if the transcriptional activity was significant (padj <0.05) and increased by at least 50 % (≥1.5 fold-change) when compared to their corresponding barcode count in the plasmid controls. Based on these criteria, 10.6% of CHD-risk variants (574 variants) exhibited CRE activity, which we refer to as “expression modulating variants” (emVars) and “expression modulating alleles” (emAlleles), respectively (**Figure 4A**).

**Figure 4:**
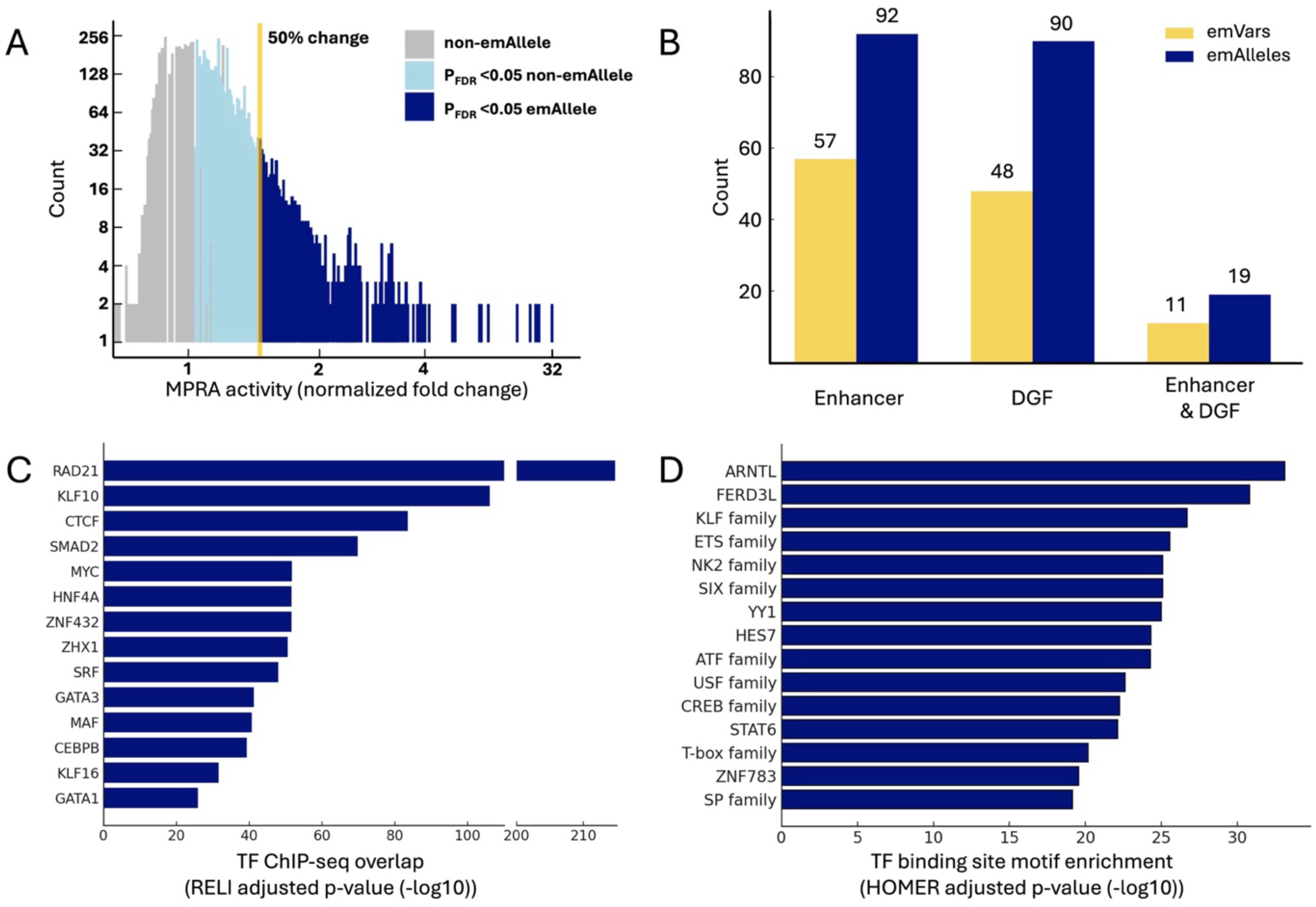
Regulatory activity of CHD-risk emVars. **A)** Distribution of MPRA regulatory activity. The normalized fold change of MPRA activity relative to plasmid control (X-axis) was calculated using DESeq2 (n = 3 biological replicates). Expression modulating alleles (emAlleles; dark blue) were identified as those alleles with significant activity relative to control (pFDR < 0.05) and at least a 50% increase in activity. Full results are provided in **Supplementary Data 3**. **B)** Overlap between emVars and cardiac regulatory elements active during heart development and the adult heart. **C)** Enrichment of regulatory protein and TFs binding at emVars. Enrichments were calculated compared to non-emVars. p-values were estimated by a one-sided z-test with Bonferroni multiple testing correction using RELI. The top 15 regulatory proteins and TFs (based on RELI p-values) that overlap at least 10% of emVars are shown. Full results are provided in **Supplementary Data 3**. **D)** TF binding site motif enrichment for emVars compared to non-emVars. p-values were estimated by one-sided hypergeometric test with Benjamini–Hochberg multiple testing correction by HOMER using the full oligo sequences of emVars and non-emVars. The top 15 enriched TF motif families are shown. Full results are provided in **Supplementary Data 3**.

We next examined the potential of the identified emVars to be meaningful in cardiovascular biology by evaluating overlap with putative enhancers and DNase I genomic footprints (DGF) active during heart development and in the adult heart. ^72,73^ Significance of the overlap between emVars and cardiac CREs was determined by a Fisher’s Exact Test using non-emAlleles (remaining oligos from the MPRA library) as a comparison (**Supplementary Figure 12**). We identified 57 emVars (∼10%; p-value = 2.10 x 10^-29^ and Odds Ratio = 4.98) that occur in cardiac enhancers, 48 emVars (∼10%; p-value = 1.57 × 10⁻^21^ and Odds Ratio = 3.73) in DGF in heart tissue, and 11 emVars (∼2%; p-value = 1.61 × 10⁻^23^ and Odds Ratio = infinite [i.e., none occurred in the non-emAllele null set]) that occur in both types of cardiac cis-regulatory elements (CREs, **Figure 4B**). From the entire MPRA library, the fact that only emVars overlapped with both types of cardiac CREs suggests a likely important role for these variants in cardiac gene regulation.

Next, we identified functional genomic features (e.g., TF and histone mark ChIP-seq peaks) enriched within emVars in cardiac CREs relative to non-emVars using the RELI algorithm. ^74^ We observed significant enrichment for ChIP-seq peaks of regulatory proteins involved in cardiovascular processes and diseases, such as RAD21 (padj = 1.59×10^-215^)^75–77^, KLF10 (padj = 8.39×10^-107^)^78–80^, SMAD2 (padj = 1.60×10^−70^)^81^, SRF (padj = 1.11×10^−48^)^82,83^, and members of the GATA family (padj < 1.11×10^−26^)^84–86^. In particular, RAD21, SRF, GATA3, and GATA1 were among the top 15 peak overlaps and have been previously implicated in the development of CHDs (**Figure 4C**). ^87–89^ Additionally, we evaluated TF motif enrichment for emVars in cardiac CREs using HOMER. ^90^ Through this analysis, we observed enrichment of multiple TF families known to play a role in heart development and cardiac diseases, such as KLF (padj <1.0×10^−11^)^91,92^, ETS (padj < 1.0×10^−11^)^44^, NK2 (padj < 1.0×10^−10^)^22,93^, CREB (padj < 1.0×10^−9^)^94,95^, and T-box (padj < 1.0×10^−8^)^17,41,42,96^ (**Figure 4D**). Many of the TF families with enriched motifs in emVars also share enriched ChIP-seq peaks from our RELI analysis (**Supplementary Data 3**). Collectively, these analyses identified CHD-risk variants that can drive gene expression from cardiac CREs, which are enriched for ChIP-seq peaks and motifs for TFs with known roles in cardiovascular development and disease progression.

### MPRA identifies 187 CHD-risk variants with allelic CRE activity

We next identified CHD-risk variants that exhibited allelic CRE activity, henceforth referred to as “allelic emVars”. Variants were considered to have allelic CRE activity if at least one allele was an emAllele, there was a significant change in CRE activity (padj <0.05), and if there was at least a 25% change in expression compared to the reference allele. Following these criteria, we identified 187 CHD-risk variants (33% of emVars, 3.4% of all CHD-risk variants) as allelic emVars (**Figure 5A**). Of the 187 allelic emVars, 169 (90.4%) had an increased CRE activity compared to their corresponding reference allele, while only 18 (9.6%) resulted in decreased activity. We then intersected the allelic emVars with the putative cardiac CREs to identify potential functional variants in cardiovascular genomics. From the 187 allelic emVars, 19 (10.2%) overlapped with cardiac enhancers, and 7 (3.7%) overlapped with heart DGFs (**Figure 5B**). From the allelic emVars, rs559405101 C>T was the only one to occur in a genomic sequence that is both a cardiac enhancer and DGF, with a 1.94-fold increase (rank 29^th^ in allelic emVars fold-change) in transcriptional activity (**Figure 5B-C**).

**Figure 5:**
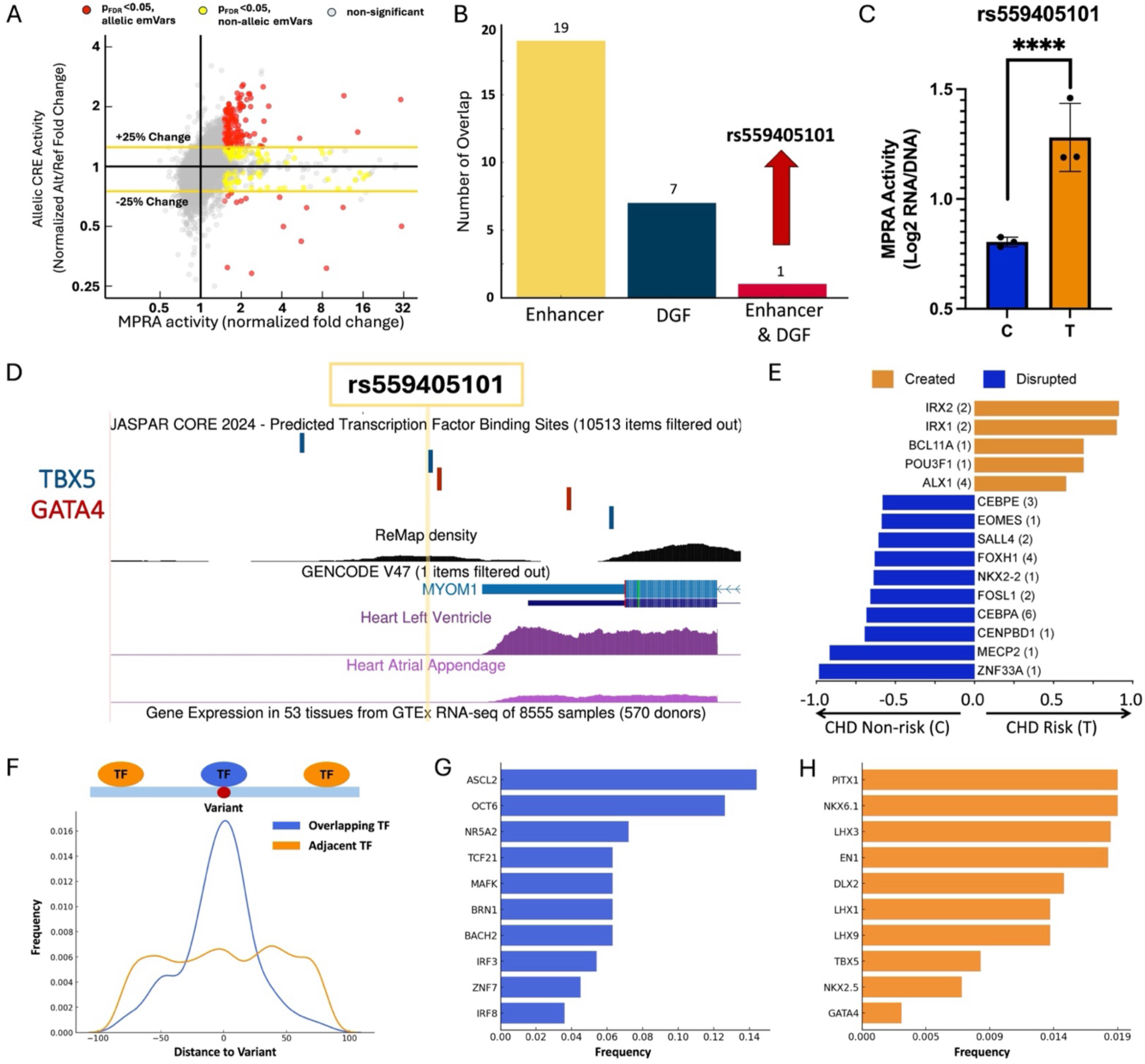
Regulatory activity and mechanisms of allelic emVars. **A)** Identification of variants with allelic CRE activity. Allelic CRE activity (Y-axis) is defined as the normalized fold change of MPRA activity between the non-reference and reference alleles (n = 3 biological replicates). MPRA activity (X-axis) is the normalized fold change of MPRA activity for any allele of the variant. Allelic emVars (red) were defined as variants with a significant difference in MPRA activity (pFDR < 0.05) between any pair of alleles and at least a 25% change in activity difference compared to the reference allele. Full results are provided in **Supplementary Data 4**. **B)** Overlap between allelic emVars and cardiac regulatory elements active during heart development and the adult heart. **C)** Normalized MPRA CRE activity of each experimental replicate for rs559405101. **D)** Genome browser map of a 2 kb window centered on rs559405101. Binding sites for GATA4 and TBX5 are displayed as blue and red rectangles, respectively. rs559405101 is upstream of *MYOM1* and is upregulated in the heart left ventricle and atrial appendage. **E)** Genotype-dependent TF binding events predicted for rs559405101. The X-axis indicates the preferred allele, along with a value indicating the strength of the allelic behavior (MARIO ARS value > 0.4), calculated as one minus the ratio of the weak to strong read counts (e.g., 0.5 indicates the strong allele has twice the reads of the weak allele). Significance (p-value < 0.05) was determined relative to binding events found in non-emVars sequences. Values in parentheses next to the TF name are the number of binding events created or disrupted by that specific TF. **F)** TF binding site location distribution for variant overlapping (blue) and variant adjacent (orange) TFs, relative to all allelic emVars. **G-H)** Motif enriched for TFs categorized as **G)** variant overlapping (Odds Ratio > 1.5, blue) and **H)** adjacent (Odds ratio < 1.5, orange) to the allelic emVars. Full results for figures 5F-H are provided in **Supplementary Data 4**.

As a possible mechanism of action, rs559405101 is flanked by three TBX5 and two GATA4 predicted binding sites, which could be key regulators of *MYOM1* in cardiac tissue (**Figure 5D**). *MYOM1* is overexpressed in muscle-skeletal and cardiac tissue (**Supplementary Figure 13**) and is known to play a role in cardiac muscle function. ^97^ Additionally, *MYOM1* has been identified as a key gene in multiple types of cardiomyopathies through cardiac gene dysregulation and alternative splicing events. ^98–100^ To further understand how rs559405101 can regulate gene expression, we identified allelic binding events of TFs (computed by MARIO using ChIP-seq data) that can potentially drive these regulatory mechanisms (**Figure 5E**). Among the top allelic binding events, we identified TFs that play established roles in cardiac development, such as IRX1/2 (heart development and function) ^101–103^ and POU3F1 (cardiac mesoderm differentiation) ^104^, which are predicted to preferentially bind to the rs559405101-T risk allele. Conversely, binding motifs of other cardiac TFs involved in heart development, such MECP2 ^105–108^ and FOXH1 ^109,110^, are disrupted by the rs559405101-T risk allele. Together, our findings suggest a possible mechanism by which the allelic emVar rs559405101 can alter *MYOM1* expression by altering biochemical interactions with multiple cardiac TFs.

Finally, we identified TFs that are predicted to have allelic binding to the remaining allelic emVars. We identified 376 unique TF motifs that only occur in either the risk or non-risk allele (**Supplementary Data 4**). However, TF-DNA interactions are not always directly disrupted by the variants, but could instead disrupt a TF binding partner or flanking region. ^15,58,111,112^ To identify which TFs tend to be variant overlapping (within 10 bps of the variant, odds ratio >1.5) versus variant adjacent (**Figure 5F**), we calculated the frequency of binding sites for each TF relative to the emVars and compared these frequencies to random expectation using a proportions test. We identified 26 high-frequency variant overlapping TFs associated with cardiac phenotypes, such as cardiac-specific immune responses (e.g., ASCL2 and IRF3) ^113–117^, cardiovascular diseases (MAFK, BACH2, and IRF8) ^118–124^, and heart function and development (OCT6 and TCF21) ^125–130^ (**Figure 5G**). Among the variant adjacent, many cardiac developmental transcription factors (TFs) were among the most frequent, such as members of the Pitx, LHX, MEF2, NK2, GATA, and T-box families (**Figure 5H**). ^96,131–136^ Altogether, integrated computational analyses identified many CHD-risk variants with genotype-dependant expression, along with regulatory genomic features and TFs that could be driving these transcriptional changes.

### CHD-risk variants exhibit allelic binding and expression

We next intersected our list of 187 allelic emVars with the 170 SNPs that exhibited allelic binding with cardiac TFs to identify potential CHD-causal variants. After overlapping the datasets, we identified five allelic emVars with allelic binding: NKX2-5 (rs2137643 and rs7303642), GATA4 (rs2896074 and rs839154), and TBX5 (rs7032149) (**Figure 6A**). The overlap between both datasets was significant by Fishers Exact Test (p-value = 0.032), suggesting that the association is likely not random. Thus, these variants were selected to be further explored as important functional CHD allelic mechanisms. The correlation between allelic binding and allelic MPRA activity throughout the CHD library was low for all three TFs (**Figure 6B**), which is not surprising considering the experimental differences between SNP Bind-n-Seq (*in vitro* with recombinant TFs) and the MPRA (cell-based assay). Additionally, less than 5% of the library actually has motifs for NKX2-5, GATA4, or TBX5. Even with motif matches, gene regulation is a highly complex process where TF binding does not always lead to transcriptional changes, as is the case with transcriptional repressor proteins and silencer elements. ^137–140^ Particularly relevant to this work, NKX2-5, GATA4, and TBX5 have all been previously reported to have dual activating and repressive functions. ^65,86,141,142^ However, variants rs2137643, rs7303642, rs2896074, and rs7032149 all resulted in a significant increase in both TF binding and expression (**Figure 6C-D**). Variant rs839154 was the only SNP to have opposite effects in regulatory mechanisms, with an increase in expression but a decrease in GATA4 binding, suggesting a repressing role for GATA4 at this locus.

**Figure 6:**
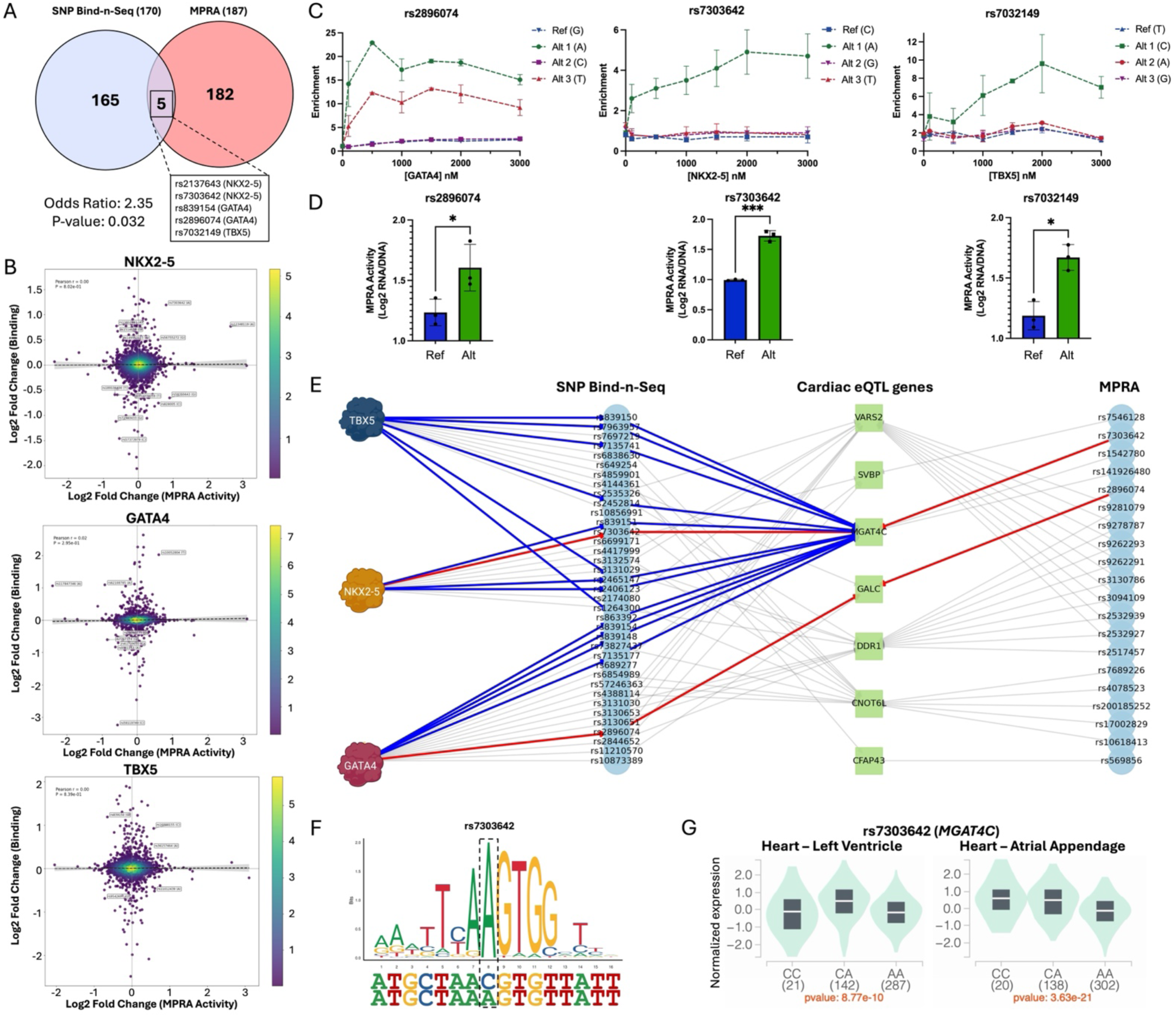
CHD-risk variants with allelic binding and regulatory activity. **A)** Venn diagram of common variants with allelic TF binding (SNP Bind-n-Seq, blue) and transcriptional activity (MPRA, red). Five common variants are displayed with the TF that showed differential binding. Significance in the association between allelic binding and gene expression was determined by Fisher’s Exact Test (p-value < 0.005). **B)** Density plot correlating expression fold change of the allelic emVars with binding fold change of NKX2-5 (left), GATA4 (middle), and TBX5 (right). Variants with a 25 % change in both binding and expression are labeled on the plot. **C)** Representative binding curve for each TF for a variant that altered the binding of NKX2-5 (top), GATA4 (middle), and TBX5 (bottom). Reference alleles (Ref) are represented in blue, and tag-SNP alleles from the GWAS catalog (Alt 1) are represented in green. Permutated alleles (alternate non-risk; Alt 2 and Alt 3) are represented in red and purple, respectively. **D)** MPRA activity for the variants used for binding curves in (**C**). **E)** Interaction networks of cardiac eQTL genes with variants exhibit allelic binding and/or gene expression. Interactions highlighted in red indicate cardiac eQTL variants that exhibited allelic behavior in binding (SNP Bind-n-Seq) and expression (MPRA). Interactions highlighted in blue indicate eQTL variants for *MGAT4C* that altered binding for all three TFs. **F)** DNA-binding motif logos are shown for NKX2-5 in the context of the DNA sequence surrounding rs7303642. **G)** eQTL indicating *MGAT4C* expression dependent on rs7303642 genotype in the heart left ventricle (data from the GTEx portal).

Next, we proceeded to identify variants with differential allelic binding (170 SNPs) and transcriptional activity (187 allelic emVars) that are eQTL in cardiac tissue (heart left ventricle and atrial appendage). From the CHD-risk variants with allelic binding, 44 (26%) were in cardiac eQTL affecting 19 genes, with over 60% sharing direction of the effect size (e.g. increase for both TF binding and eQTL effect size; **Supplementary Figure 14A**). From the allelic emVars, 34 variants were also in cardiac eQTL with 33 genes (**Supplementary Figure 14B**). We identified seven eQTL genes that overlapped between the SNP Bind-n-Seq and MPRA variants and constructed interaction networks with both sets (**Figure 6E and Supplementary Figure 14C-D**). An interaction network revealed that rs7303642 and rs2896074 were eQTL for two overlapping genes, *MGAT4C* and *GALC*, respectively. MGAT4C (Mannosyl Alpha-1,3-Glycoprotein Beta-1,4-N Acetylglucosaminyltransferase) is an enzyme that is highly expressed in the heart and has been linked to CHDs, such as conotruncal heart defects. ^143,144^ Additionally, 14 eQTL variants for *MGAT4C* altered the binding affinity of NKX2-5 (4 SNPs), GATA4 (5 SNPs), and TBX5 (5 SNPs) when biochemically evaluated through SNP Bind-n-Seq (**Figure 6F-G, Supplementary Figure 15**). GALC (Galactosylceramidase) is an enzyme involved in galactose metabolism that has been implicated in several diseases, particularly respiratory and neurological diseases. ^145,146^ Although having modest expression in cardiac tissue, to date, there is no *GALC* implication with specific cardiovascular diseases or phenotypes.

Finally, the interaction network revealed additional cardiac eQTL genes with eQTLs that displayed altered binding of all three TFs tested, such as *CNOT6L*, *VARS2*, and *DDR1*. For example, 20 SNPs that are *CNOT6L* eQTL also that exhibited allelic binding for the three TFs (**Supplementary Figure 16**). CNOT6L is a protein subunit of the CCR4-NOT complex, which regulates mRNA deadenylation and has been implicated in proper heart functioning. ^147–151^ In an interesting case, *VARS2* and *DDR1* are eQTL genes of the same eight variants with allelic binding, where *VARS2* consistently had an increased effect size and *DDR1* a decreased effect size (**Supplementary Figure 17**). *DDR1* is a discoid domain receptor that is involved in cardiovascular calcification, while *VARS2* is a tRNA synthetase that has been linked to heart failures when mutated. ^152–155^ In summary, our combined findings provide a high-confidence list of possible mechanisms of CHD-risk variants that exhibit allelic binding, transcriptional activity, and cardiac eQTL.

## Discussion

Genome-wide association studies (GWAS) have linked thousands of non-coding genomic loci to congenital heart disease (CHDs). ^51–55,156–159^ However, translating these associations into meaningful biological mechanisms remains a challenge. This work provides the first combined genome-wide evaluation of cardiac TF binding and transcriptional activity, encompassing over 3,000 CHD-risk variants. Our approach provides crucial information needed to understand biochemical and genetic mechanisms underlying CHD etiology, and ultimately for identifying causal variants.

To date, over 90% of CHD-associated variants have been mapped to the non-coding genome and thus could potentially alter the function of cardiac TFs during heart development. ^2,4,6,10^ With SNP Bind-n-Seq, we build upon previous experimental approaches ^45,46^ to assay thousands of variants in a high-throughput manner. Comparted to other techniques to profile TF binding, SNP Bind-n-Seq can quantify enrichment across all variant alleles simultaneously and determine allelic binding constants, which has previously not been done. As proof of concept, we chose NKX2-5, GATA4, and TBX5, three cardiac TFs crucial for heart development that have been implicated in CHD etiology, to evaluate CHD-risk variants for allelic binding events. ^17,18,20–22,26,66,160^ Using SNP Bind-n-Seq, we constructed allelic binding curves for 3,232 CHD-risk variants (12,928 alleles) for all three cardiac TFs. Our findings provide ∼38,400 unique DNA binding affinity measurements, with 5.3% of variants exhibiting allelic binding for at least one of the three TFs tested in this study. This approach also provided mechanistic insight into the observed allelic binding events, with only 74% of the examined SNPs directly creating or disrupting the binding motifs of NKX2-5, GATA4, and TBX5. The remaining 26% altered cardiac TF binding by creating low-affinity binding sites adjacent to core binding motifs or occurred within TF binding flanking regions.

The data generated through SNP Bind-n-Seq were subsequently used to train computational models to predict changes in TF-DNA binding. In doing so, we scored all of the disease-associated variants contained in the GWAS catalog to predict allelic binding events for NKX2-5, GATA4, and TBX5. We identified over 1,000 variants predicted to alter the binding of these three TFs that are associated with multiple types of diseases, not just cardiovascular.. Thus, this study provides insight into potential roles for NKX2-5, GATA4, and TBX5 in disease etiology outside of CHDs. SNP Bind-n-Seq is a scalable approach that can be implemented with other TFs or DNA-binding proteins to study biochemical mechanisms of multiple genetic diseases associated with non-coding variants.

We complemented our *in vitro* binding affinity data with transcriptional activity in a cellular context using an MPRA to quantify changes in transcriptional activity. Since our previous approach was based on three cardiac TFs involved in heart development, we performed the MPRA in a Flp-In 293 system that was genetically modified to stably express NKX2-5, GATA4, and TBX5. Our results showed that 574 emVars (10.6% of tested variants) drove CRE activity, and of those, 16.4% occur within known cardiac CREs that are active during heart development. ^72,73^ We identified 187 CHD-risk variants (3.5% of tested variants) with allelic CRE activity, of which 13% occurred in cardiac CREs. Our downstream computational analyses also provide mechanistic insight into the likely regulatory proteins driving these transcriptional changes, along with their affected target genes. Our MPRA findings have some limitations, including the cell line of choice, which lacks cardiac-specific chromatin features and other cardiac TFs beyond those within the scope of this work. However, previous studies on the genotype dependence of cis-eQTLS have shown that variants exhibit a bimodal distribution, where expression patterns are either single tissue-specific or shared across most tissues and cell types. ^161–163^ Additionally, previous work has shown that MPRA activity is highly correlated across multiple cell types. ^164^

Finally, our study reveals possible mechanisms of CHD-risk variant that may make important functional contributions to CHD. After combining our findings from the SNP Bind-n-Seq and MPRA assays, we identified five variants (rs2137643, rs7303642, rs2896074, rs839154, and rs7032149) that can be further studied for important roles in CHD. Notably, variants rs7303642, rs2896074, and rs839154 are eQTLs for genes expressed in cardiac tissue, which may further support their role in cardiovascular disease genomics. Additionally, the combined findings identified variants with allelic regulatory activity that were cardiac eQTL for seven genes. Among them, *MGAT4C*, *CNOT6L*, *VARS2*, and *DDR1* were the eQTL genes of multiple variants exhibiting allelic binding for NKX2-5, GATA4, and TBX5. Altogether, our combined findings through high-throughput TF-DNA binding and reporter assays provide the largest biochemical and genetic evaluation of thousands of CHD-risk variants.

With the advancement in sequencing technologies and rapid identification of variants through GWASs, identifying and validating disease-causing variants remains challenging. In this study, we present a functional framework for identifying and assessing thousands of variants that alter regulatory mechanisms, including TF binding and regulatory activity. We believe that SNP Bind-n-Seq, coupled with other high-throughput methods like MPRA, is a scalable approach to dissect the regulatory mechanisms of non-coding variants and understand the genetic etiology of many complex human diseases.

### Limitations of Study

This work, although providing valuable insight into biochemical and genetic mechanism behind CHD-risk variants, has limitations that can be addressed in future experiments. First, *in vitro* binding assays were performed on purified TF DBDs from a bacterial expression system. This has limitations, such as proteins lacking post-translational modifications (PTMs) that occur in eukaryotic cells. Since binding is performed *in vitro*, biochemical interactions are performed out of cellular context without binding partners and co-factors. Additionally, binding is performed using short (40 bp) synthetic DNA sequences lacking sufficient genomic context. Second, the reporter assay was performed on genetically modified Flp-In 293 cell lines to express the three cardiac TF from the binding assays. This system lacks the cardiac-specific cellular context and gene expression profiles of biologically relevant cell lines, such as cardiomyocytes and cardiac progenitors. In the future, our framework can be adapted to perform binding assays using nuclear extracts of cardiac cell lines, as well as to conduct the MPRA at multiple developmental stages of cardiomyocyte differentiation.

## Materials availability

The CHD libraries used for the SNP Bind-n-Seq and MPRA, and the genetically modified cell lines used in this work, are available upon request.

## Data and code availability

The original code designed in this work for SNP Bind-n-Seq is available at: https://github.com/Shreya-droid/SNPoiss_bind_n_seq. Code for MPRA barcode count and analysis is available at: https://github.com/tewhey-lab/MPRAmodel. Code for MPRA plotting is available at: https://github.com/WeirauchLab/mpraprofiler. Code for RELI analysis is available at: https://github.com/WeirauchLab/RELI.

## Author Contributions

EGPM and JARM designed and conceptualized the project. EGPM, DAPM, JLMB, LSA, ARM, and RVR performed protein purifications and TF-DNA binding assays (SNP Bind-n-Seq and validations through EMSA). SS, JGMF, JAFR, and DB performed computational analysis for the SNP Bind-n-Seq sequencing data. EGPM, ACBR, JMRR, HH, KD, and MTH performed Flp-In 293 genetic modification and cell culture. EGPM, ER, LGP, and MG performed MPRA experiments. EGPM, MN, OAD, ER, and XC performed computational analysis of MPRA data. EGPM and SS performed plotting and data visualization. LPL, CF, MTW, LCK, SKR, DB, and JARM supervised the work, provided mentoring, and secured funding for the project. EGPM, SS, DB, and JARM wrote and reviewed the original manuscript. All authors read, assisted in editing, and approved the final version of the manuscript.

## Acknowledgments

This project was supported by NIH-SC1GM127231, NSF 1736026, University of Puerto Rico Rio Piedras Institutional Funds (FIPI), Puerto Rico Science, Technology, and Research Trust, and an NIH Institutional Development Award (IDeA) INBRE (P20GM103475W). EGPM, JMRR, RVR, DAPM, ACBR, LSA, and NEMP were funded by the NIH RISE Fellowship (5R25GM061151–20). JLMB and ARM were funded by the NSF PR-LSAMP fellowship (HRD-2008186). DAPM was funded by NSF [IQ BIOREU 1852259]. EGPM and JMRR were funded by the NSF BioXFEL Fellowship (STC-1231306). ARM was funded by the NSF REU funded ARM: PR-CLIMB Program (2050493) and NIH 1T34GM145404. LSA was funded by the NIH ID-GENE Fellowship (1R25HG012702–01). JMRR was funded by an NSF graduate research fellowship (1744619). MN was supported by a Gruber fellowship. MTW and LCK were funded by NIH grants R01NS099068, R01AI024717, R01AI148276, U24 HG013078, U01 HG011172, and P30AR070549. SKR was funded by R01HG012872. The graphical abstract and Figure 1A were made using BioRender.

## Materials and Methods

### Variant selection and DNA sequence generation

For the SNP Bind-n-seq experiment, we downloaded 121 CHD-associated SNPs from six studies in several populations from the GWAS from the catalog. ^51–56^ Variants in linkage disequilibrium (LD) from four populations (EUR, AFR, EAS, SAS, R^2^=0.80) were included using the TOP LD web tool. ^165,166^ Insertions, deletions, and incomplete entries from the GWAS catalog were removed. All variants were permutated to include every possible nucleotide for each CHD-associated variant. We retrieved 40 bps of hg38-flanking DNA sequences for every allele, with the variant located in the center (19 bps upstream and 20 bps downstream of the variant). Adapters and unique molecular identifiers (UMIs) were added to each sequence at either end (5′-TCCCTACACGACGCTCTTCCGATCT – NN - [40 bp oligo] - NN -GATCGGAAGAGCACACGTCTGAACTCCAGTCAC −3′) to make a 102 bp DNA sequence. A total of 13,039 oligos (3,232 variants, 12,928 alleles, 91 cardiac DGF, and five control variants) were obtained from Custom Array.

For the MPRA, we downloaded 157 CHD-associated SNPs from the nine GWAS in several populations that were expended for variants in linkage disequilibrium (LD, R^2^> 0.8) based on 1000 Genomes Data in the ancestry(ies) of the initial genetic association using PLINK(v1.90b) as previously described. ^47,167^ All expanded variants were updated to the dbSNP 155 table from the UCSC table browser based on either variant name or genomic location. Unmappable variants were discarded. For single nucleotide polymorphisms, we pulled 170 base pairs (bps) of hg38-flanking DNA sequences for every allele, with the variant located in the center (84 bps upstream and 85 bps downstream of the variant). For the other types of variants (indels), we designed the flanking sequences to ensure that the longest allele has 170 bps. Adapters (15 bps) were added to each sequence at either end (5′-ACTGGCCGCTTGACG - [170 bp oligo] - CACTGCGGCTCCTGC-3′) to make a 200 bp DNA sequence. For all resulting sequences, we created a forward and reverse complement sequence to compensate for possible DNA synthesis errors. A total of 29,048 oligos (5,431 variants, 14,524 alleles) were obtained from Twist Bioscience.

### NKX2-5, TBX5, and GATA4 expression and purification

The human NKX2-5 homeodomain (HD) coding sequence (Asp16 to Leu96) was cloned in pET-51(+) expression vector containing an N-terminal Strep•Tag II® and a C-terminal 10× His•Tag® through Gibson Cloning and purified through Ni-NTA affinity chromatography, as previously described. ^35^ The coding sequence for the human TBX5 T-box domain (Met51 to Ser248) was cloned into a pET expression vector with a 6x His•Tag® (VectorBuilder Inc.) and purified through Ni-NTA affinity chromatography, as previously described. ^35^ Human GATA4 DNA-binding domain (DBD), also known as a zinc finger domain (ZF), gene (Uniprot: P43694, Met207 to Ala333) was cloned into the pET-28a(+) vector with an N-terminal 6x-Histag (Twist Bioscience).

### SNP Bind-n-Seq

An oligopool of CHD-associated variants (Custom Array, Supplementary File) was amplified for 12 cycles by touchdown PCR (denaturing at 95 °C for 20 sec, annealing at 70 - 0.5 °C/cycle for 15 sec, and extension at 72 °C for 15 sec) using KAPA HiFi HotStart ReadyMix (Roche Sequencing Store, #KK2602 07958935001). Sequences were amplified with IR700 fluorescent and biotinylated primers for subsequent gel excision and extraction, respectively.

Binding reactions for TBX5 and NKX2-5 were 20 µL (50 mM NaCl, 10 mM Tris-HCl (pH 8.0), 10% glycerol, 1000 ng pdI-dC, 0.5 µg BSA, 0.02% Tween-20, and 1 mM DTT) containing 50 ng of oligopool library DNA. For GATA4, binding reactions were 20 µL (10 mM HEPES (pH 8.0), 0.5 mM ZnA, 100 mM NaCl, 5% Glycerol, 1000 ng pdI-dC, 0.5 µg BSA, 0.02% Tween-20, and 1 mM DTT) containing 50 ng of oligopool library DNA. For each TF, six concentration points were used for binding assay: 0, 100, 500, 1000, 1500, 2000, and 3000 nM. Reactions were incubated for 30 min at 30 °C, followed by a 30 min incubation at room temperature, and samples were loaded onto a 6% polyacrylamide gel in 0.5x TBE (89mM Tris/89mM boric acid/ 2mM EDTA, pH 8.4). The gel was pre-ran at 70 V for 15 min, loaded at 30 V, and resolved at 120 V for 2 hours at 4 °C. The gels were revealed using Azure Sapphire Bio-molecular Imager with 658nm/ 710nm excitation and emission.

After EMSA, bound and unbound bands for each concentration point were cut, and DNA was extracted and left overnight in 500 µL of EB buffer (Qiagen, #19086) at 30 °C shaking at 1200 rpm (Thermoshaker, BIOGRANT). After gel extraction, DNA was purified using Dynabeads M-280 Streptavidin (Invitrogen, #11205D) according to manufacturer procedures. Barcodes and sequencing adapters were added to bound and unbound sequences through 13 cycles of PCR (denaturing at 95 °C for 30 sec, annealing at 64 °C for 30 sec, and extension at 72 °C for 30 sec).

### SNP Bind-n-Seq datasets and preprocessing

The sequencing was performed using the Illumina platform, generating paired-end reads of 150 base pairs (PE150). This produces two reads per DNA fragment one from each end. For this analysis, only Read 1 (from the 1.fastq file) was used. Next, to extract our genomic fragment which is less than 150bp, we used the constant right flanking region from the library structure “GATCGGAAGAGCACACGTCTGAACTCCAGTCAC” to locate 44 bp DNA sequence that includes a SNP position. This is done with all the FASTQ files for each TF: NKX2-5, GATA4, and TBX5. Our library contained a 40 bp genomic DNA sequence along with a 4 bp Unique Molecular Identifier (UMI), two at each end. For each sequencing read, we generated a pattern where the SNP at the 20th position could be occupied by any of the four nucleotides: A, T, G, or C. We then calculated the frequency of each 4 possible nucleotide at the SNP position across all sequencing reads, giving us the counts for each SNP position for a sequence.

### Enrichment Analysis

For each nucleotide at a given SNP position, enrichment was calculated relative to the unbound dataset. To avoid division by zero and reduce the impact of low-count noise, pseudo-counts were added to both bound and unbound read counts. The pseudo-counts were defined as a small background signal equivalent to 2.5 reads per million, scaled by the total number of reads in the respective bound or unbound dataset. Quantitatively, the pseudo-counts were computed as (2.5 X total reads)/10^6^. Enrichment for each nucleotide was then calculated by dividing the pseudo-count-adjusted count in the bound sample by the corresponding adjusted count in the unbound sample. This approach allows a robust comparison between bound and unbound conditions, generating enrichment profiles that reflect nucleotide-level binding preferences at SNP positions.

Before calculating the enrichment profile, we ensure data quality and consistency, and thus remove low-quality sequences based on specific thresholds designed to minimize deviations that could compromise data uniformity. To control for differences in sequencing depth between samples, the raw counts of each unique sequence, regardless of the nucleotide identity at the SNP position, were normalized by the total number of sequences in the corresponding sample. These normalized values were then scaled to parts per million (PPM). Then, the thresholds applied were as follows:

1. **Filtering Sequences from 0nM Unbound Condition**: To ensure a better analysis, a ratio threshold of **≤ 0.2 or ≥ 2** was applied to the 0nM Unbound-to-Library ratio.

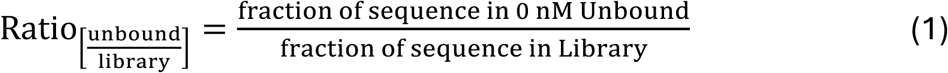 Samples with a 0 nM Unbound-to-Library ratio outside the ≤ 0.2 or ≥ 2 range were excluded from further analysis to minimize bias from gel-based irregularities.
2. **Low Counts in 0nM Unbound**: Sequences with read counts **less than 5 per million** (in both replicates, R1 and R2) in the Unbound fraction at 0nM were excluded, as this low representation indicates a lack of binding specificity at baseline.
3. **Low Counts in Library**: Sequences with read counts **less than 5 per million** in the original library sample were also removed to ensure that subsequent analyses focused only on sequences present in suÜicient abundance.
4. **Low Ratio Threshold for 0nM Bound-to-Unbound Analysis**: A ratio threshold of **≤ 0.2 or ≥ 5** was applied to identify sequences highly enriched in the Unbound fraction (indicating low binding aÜinity) or overly represented in the Bound fraction (suggesting nonspecific binding at 0 nM).
5. **Sucicient Representation of Nucleotides**: Sequences with counts **below 5 per million** across all nucleotide bases (A/T/G/C) were excluded to ensure analyses included only sequences present in adequate quantities across all samples.

These criteria were essential for refining the dataset, focusing on sequences with meaningful binding behavior while eliminating potential artifacts or non-specific interactions. For those SNP sequences that have passed these thresholds, then the enrichment calculation can be expressed as the ratio of the relative frequencies of a base in the sample data to its unbound sample. The Enrichment is defined as the ratio of relative fractions as following:

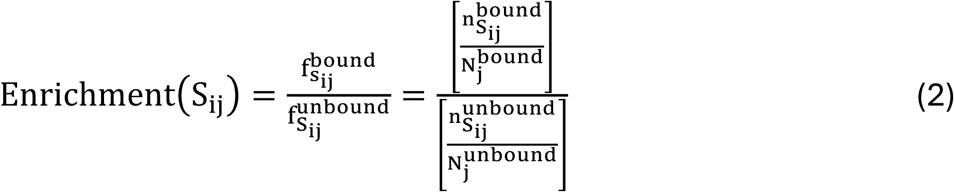

where,

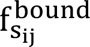 fraction of sequence S_i_ at j^th^ concentration in the bound sample,

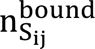 is the count of sequence S_i_ at j^th^ concentration in the bound sample,

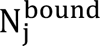 is the total sequence count at the j^th^ concentration in the bound sample,

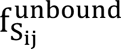 fraction of sequence S_i_ at j^th^ concentration in the unbound sample,

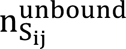 is the count of sequence S_i_ at j^th^ concentration in the unbound sample,

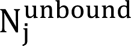 is the total sequence count at the j^th^ concentration in the unbound sample.

Enrichment indicates how much more (or less) frequent a sequence is occurring in the sample compared to its corresponding unbound sample. The significance of these calculations is that the enrichment value gives you an idea of how over- or under-represented a sequence is in your data compared to an unbound fraction. Enrichment was calculated for every sequence for each factor at all concentrations and for both the replicates, this was then used for plotting. After applying all the above thresholds, we generated a dataset that contains binary values (0s and 1s) for each SNP sequence (rsID’s), indicating whether the respective sequence passed or failed each of the specified thresholds (see section). Only those rsID’s that met all threshold criteria were retained for subsequent analyses, including model fitting.

### Estimating Binding Affinity and Probability of Molecular Interaction

The probability of TF binding to DNA depends on the sequence-specific binding affinity, driven by direct molecular contacts like hydrogen bonds and hydrophobic interactions between TF amino acids and DNA bases. This affinity is quantified by the dissociation constant *K_d_*, which reflects the free energy of binding. TF concentration influences binding via mass action, while chromatin accessibility modulated by nucleosome positioning and epigenetic modifications regulates physical DNA availability. Nonspecific binding results from weaker electrostatic interactions with the DNA phosphate backbone and bases outside consensus motifs, contributing to TF search efficiency and buffering TF concentration.

Here, we use a statistical mechanics approach to derive binding affinity and other equilibrium properties of the system. In our experiment the protein is exposed to a pool of 12940 SNP sequences simultaneously, each sequence consists of 44 nucleotides containing up to *n* number of cognate binding sites. Assuming independent binding at each site the occupancy probability of sequence *S_i_* at concentration *j* is calculated as

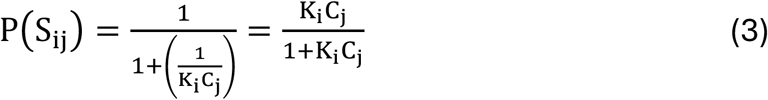

where, K_i_ is the association constant for sequence i at the j^th^ concentration, C_j_.

To account for DNA outside bound and bound fraction (e.g., stuck in the well of the gel, uneven gel migration, etc.), we introduced an additional parameter *w_i_*, representing the probability that a given DNA molecule is *“stuck in the well”.* This decision further supported by insights from the enrichment line plots, which revealed that previously fitted K-values did not fully capture the behaviour of certain sequences. These deviations prompted us to introduce an additional parameter, *w_i_*, to better model such effects and more accurately represent the binding behaviour under these conditions.

To substantiate the use of an additional parameter, *w_i_*, we examine read count distributions across three experimental fractions: the original library (L), the unbound fraction (U), and the bound fraction (B). The library contains the initial representation of all DNA sequences before exposure to the TF, while the unbound fraction contains sequences that did not bind to the TF, and the bound fraction contains those that did. At 0nM TF, where no specific TF-DNA interactions should occur, we expect most sequences to remain in the unbound fraction. That is, for any sequence *i*, its unbound read count (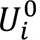) fraction should be approximately equal to its initial library count (*L_i_*) fraction:

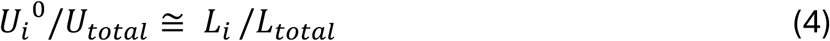

However, when *U_i_*^0^/*U_total_* ≅ *L_i_* /*L_total_*, indicates that sequence *i* is unexpectedly depleted from the unbound fraction, even though no TF is present to cause this. This suggests the sequence might be subject to non-specific loss, perhaps binding to the experimental surface, interacting with background proteins, or being inefficiently recovered. More generally, if we observe that the total recovered reads *B_i_*/*B_total_* + *U_i_*/*U_total_*, are much smaller than the initial library pool (*L_i_*), i.e.,

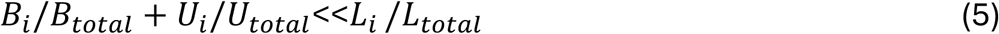

This supports the same conclusion that the sequence is systematically underrepresented and should be excluded to avoid introducing artifacts into the binding analysis.

In the updated equation we defined the range of *w_i_* ∈ [0,1], where *w_i_* = 0 means the molecule is entirely unbound or completely stuck in the well and *w_i_* = 1 completely bound with the TF, suggesting strong binding. Thus, we updated the probability of sequence to be found in the bound region of gel is given by equation (4)

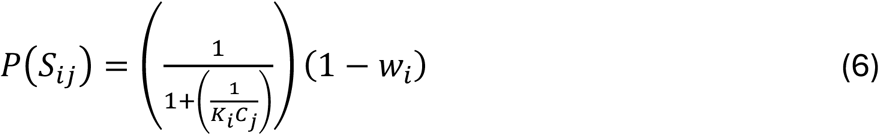

Our modelling approach integrates bound and unbound read counts along with library data to jointly infer the key kinetic parameters. Due to the competitive binding environment, these constants must be inferred jointly, as the observed binding reflects the relative affinities of all sequences in the pool. The model specifically optimizes the concentration-independent parameter *K_i_*, which controls the observed SNP read counts across varying protein concentrations, thereby capturing the system’s dynamic behaviour. In this framework:

Using the Poisson probability mass function:

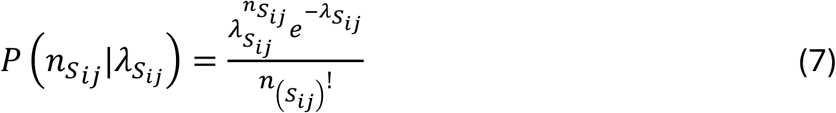

given the observed counts *n_sij_*, assumed to follow a Poisson distribution

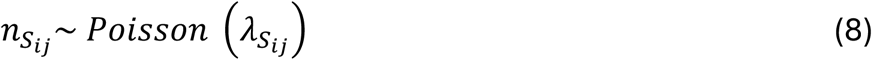

In our model, each sequence i is associated with a kinetic parameter *K_i_*, which represents its effective association constant and determines its binding affinity in the competitive environment. The expected read count *λ_s_ij__* for sequence *i* at *j* is a function of *K_i_* and the protein concentration, capturing the binding probability under equilibrium assumptions. Then the likelihood function is

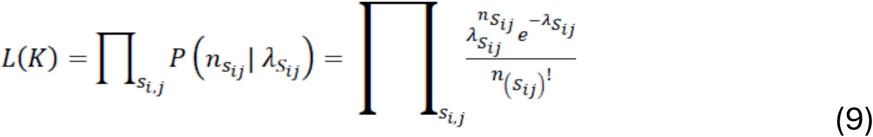

Taking the logarithm, log-likelihood function is:

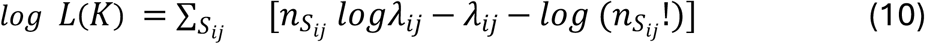

The goal is to maximise the concentration independent parameter K, which influences the likelihood of observing read counts for SNPs at different concentrations.

### Modelling the Expected Count *λ_s_ij__*

The expected fraction of sequences *S_i_* in the bound at concentration *C_j_* can be estimated as:

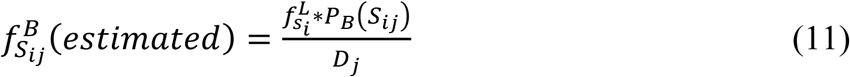

where,

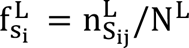 is the fraction of sequence S_i_ at j^th^ concentration in the library,

*P_B_*(*S_ij_*) = probability of bound sequence *S_i_* at concentration *j*,

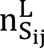 is the count of sequence S_i_ at j^th^ concentration in the library,

*N_L_* is the total sequence count in the library,

*D_j_* is the denominator at *j^th^*concentration in our enrichment equation. This is the sum of all weighted binding values over both replicates (see eq. 6). The denominator, *D_j_* accounts for the total expected binding signal from all variants in the library at *j^th^*concentration, this is critical for normalization. Initially these denominators were initialised with the (0.9/maximum enrichment), then a Poisson model without *w_i_* parameter ran up to 1000 iterations. After *360^th^*, we observed these denominators converge. So, we use 361^st^ denominator values for the model equation. We can also write as:

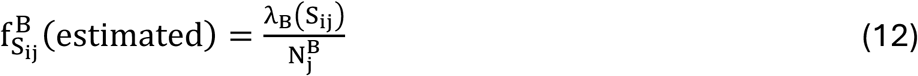

where λ_B_(S_ij_) is the expected counts of sequence S_i_ in the bound fraction at the j^th^ concentration, C_j_. Therefore,

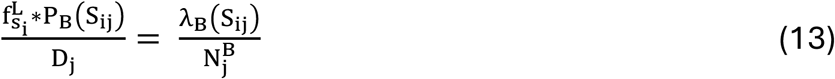

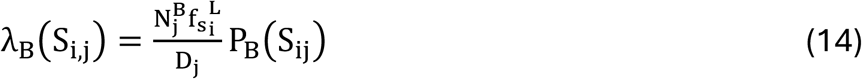

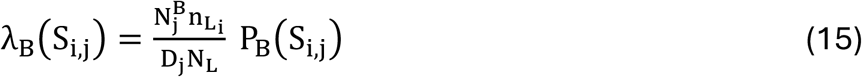

Using equation (4) we get:

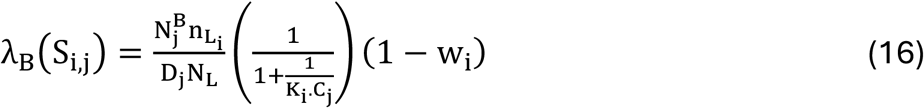

Expected count estimate of sequences S_i_ at concentration C_j_ in case of unbound:

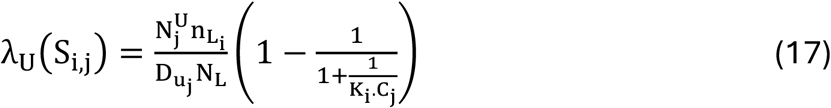

where, 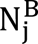 and 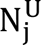 are total sequence count at the j^th^ concentration in the bound and unbound sample respectively. D_j_ and D_uj_ are the sum of all weighted binding values over both replicates in the bound and unbound sample respectively.

The total log-likelihood, as given in Equation (10), is expressed as:

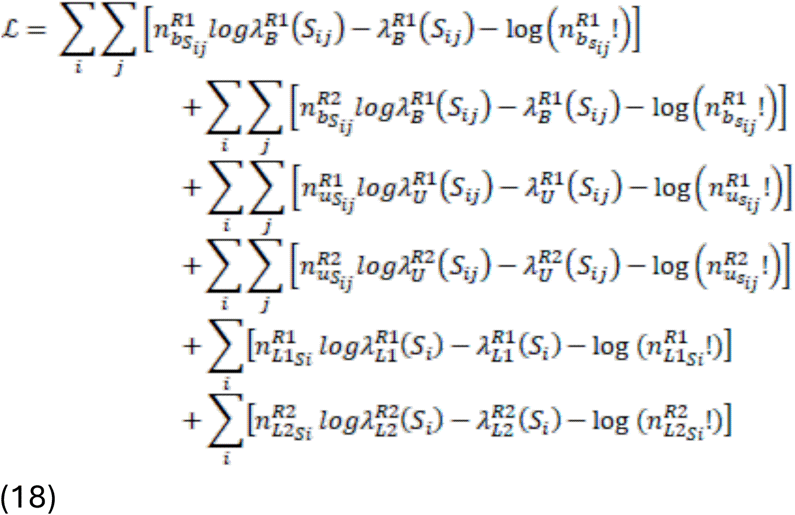

where, 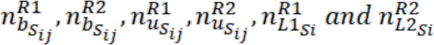 are the count of sequence *S_i_* at *j^th^* concentration in the bound (b), unbound (u) and library (L) for replicate 1 (R1) and replicate 2 (R2). Note: 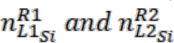 are not indexed by concentration, as the input library is shared across all concentrations. 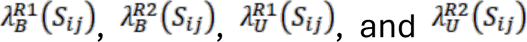 represent the expected counts of sequence *S_i_* in the bound (B) and unbound (U) respectively, for replicate 1 (R1) and replicate 2 (R2) at *j^th^* concentration, as defined in equations (16) and (17). In contrast, 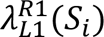 and 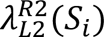 denote the expected counts of *S_i_* in the input library (L) for R1 and R2, which are independent of concentration.

The term 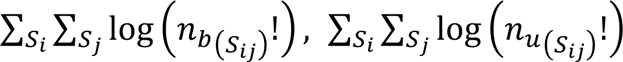, and 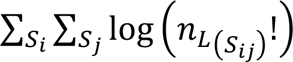, depends only on the observed data and not on the parameter*λ*. We therefore collect these data-dependent terms and denote them as a constant *C* :

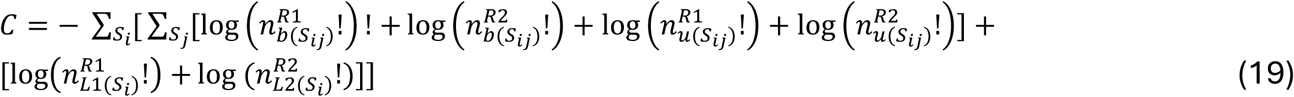

Since the constant term *C* does not depend on the model parameters, we separate it from the full log-likelihood expression. The optimization is then performed by minimizing the negative log-likelihood excluding *C*, as it has no effect on the parameter estimates. Upon eliminating the constant term from the right-hand side, the equation reduces to:

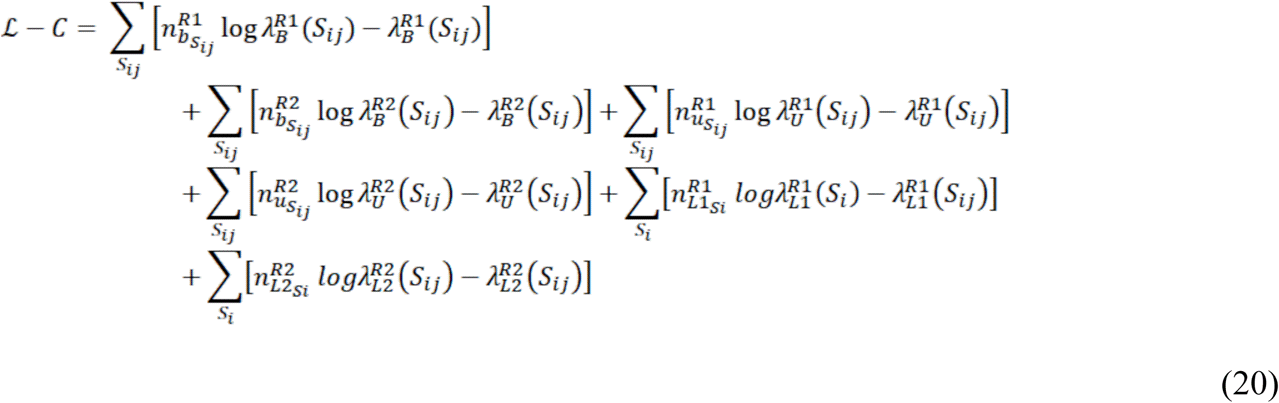

We minimize the negative log-likelihood to assess how well the model fits the observed data; this optimization simultaneously addresses a mathematical problem and holds biological significance in our context. Since optimizing all parameters at once is highly time-consuming, we instead adopt an iterative refinement procedure. To estimate parameters that minimize the negative log-likelihood, we used the *“minimize”* function from Python’s *scipy.optimize* module. The model fitting proceeds iteratively to update denominators and parameters. First, in the iteration step: for each SNP sequence, we optimized parameters by minimizing the negative log-likelihood. Then using the fitted parameters, we updated the denominators to better reflect the current binding landscape. Parameter estimation was performed through 10 iterations of optimization. Starting with initial parameter guesses, likelihoods were computed from observed count data. Using the L-BFGS-B method, a quasi-Newton method that is efficient for large scale problems and capable of handling bound constraints, the model parameters were iteratively updated to maximize likelihood. At each iteration step, the updated values of *K*, *λ*, and *w_i_* were logged. Normalization terms were also updated accordingly. This iterative procedure progressively refined the estimates and enabled effective tracking of convergence across iterations.

We started this modelling approach to better understand the relationship between TF binding and concentration. By analyzing and maximizing the fit to the observed data, we aim to accurately capture the biological patterns underlying binding behavior. This is especially important because plotting enrichment values across different concentrations and between replicates reveals variability that is non-trivial, motivating a quantitative and rigorous model.

Note: The 500 nM concentration for TBX5 was excluded from binding curve fitting and subsequent plotting due to insufficient counts in the bound fraction.

### MEME motif discovery

Upon initial receipt of the raw FASTQ files, a preliminary analysis was conducted to verify the presence of TF binding motifs within the sequencing data. To this end, we ran MEME on the complete dataset encompassing all three TFs: NKX2-5, GATA4, and TBX5. This analysis successfully identified the canonical motifs corresponding to each TF, confirming the validity and quality of the sequence data.

To further characterize motifs in individual samples relative to concentration, a more selective approach was used. For each TF, we selected the top 500 sequences with the highest enrichment scores across all four SNP variants (A, T, G, C). These subsets were then re-analysed using MEME to identify the most representative motifs under conditions of strong signal. MEME processes a group of sequences and outputs the requested number of motifs, using statistical modelling to automatically select the best motif width, number of occurrences, and description. To constrain the search and improve motif detection, specific flags were applied during MEME runs: the ***-meme-mod*** *anr* flag assumes each sequence can contain any number of non-overlapping occurrences of each motif, which is useful for detecting multiple repeats within a sequence and provides greater accuracy but requires about ten times more computational time and is less sensitive to weak non-repetitive motifs; the ***-minw*** *5* and ***-maxw*** *9* flags limit motif lengths to between 5 and 9 bases, focusing on short motifs; and the ***-meme-nmotifs*** *5* flag restricts the search to up to five distinct motifs. Together, these settings control the motif discovery process to identify a specific number of short, potentially repetitive patterns within the selected sequences.

### Motif analysis by SNP position

To investigate SNP-driven disruptions in TF binding motifs, sequence extraction was initiated using the top 500 K-fitted values. These sequences served as input for the MEME suite, executed in “anr” mode with motif width parameters set between 5 and 6 nucleotides for NKX2-5. The analysis permitted the identification of up to 10 motif logos, considering both the forward and reverse complement strands. Among the identified motif logos, the highest scoring Position Weight Matrix (PWM), regardless of strand orientation, was selected for downstream analysis.

The SNP motif analysis aimed to quantify the impact of variants on TF binding, specifically assessing changes induced by alternate alleles. For each SNP sequence, unique molecular identifiers (UMIs) were removed from both sequence termini, and 40-nucleotide windows were constructed to incorporate either the reference or alternate allele. Reverse complement sequences were also generated. These sequences were scored using the selected PWM to assess motif affinity, and the highest-scoring subregions were identified for each case. SNPs were evaluated for their potential to disrupt or enhance transcription factor binding by examining whether the SNP position overlapped with the most affected k-mers. Affected and unaffected motifs were quantified separately for SNPs associated with increased or decreased binding affinity, enabling comparative analyses across conditions.

To visualize the positional influence of SNPs within motifs, bar plots were generated to depict the frequency of SNP overlap at each motif position, stratified by directionality of enrichment. Additionally, the relative contributions of SNPs to individual motif positions were assessed by aggregating the maximum scoring bins across increased and decreased binding categories. These contributions were further visualized using customized bar plots, with specific bin ranges (e.g., positions 16–20) renamed for interpretability (e.g., positions 1–5), facilitating focused inspection of key regions. This motif disruption analysis was conducted for the transcription factors GATA4, TBX5, and NKX2-5, providing detailed insights into the allelic effects of SNPs on transcription factor binding across these critical cardiac regulators.

### Model training and testing

To evaluate the predictive performance of various models, we selected the top 500 SNPs based on their K-values. These SNPs were randomly shuffled and subsequently divided into 300 SNPs for training and 200 SNPs for testing. In parallel, we utilized ChIP-seq datasets for the cardiac transcription factors NKX2.5 (SRX9284027), GATA4 (SRX9284038), and TBX5 (SRX2023721). From each ChIP-seq dataset, the top 1000 genomic sequences were selected based on peak signal intensity scores and were similarly partitioned into training and testing sets.

We evaluated four cases:

1. **SNP-based model**: A classification model was trained using 300 positive and 300 negative SNP sequences, and evaluated on the remaining 200 positives and 200 negatives from a total of 500.
2. **ChIP-based model**: A model was trained using 600 positive and 600 negative ChIP sequences, and evaluated on the 400 positives and 400 negatives.
3. **ChIP-seq-trained model, SNP-seq tested**: A separate model was trained using 1000 ChIP-seq peak sequences (1000 positive and 1000 negative sequences) and tested on the 500 SNP dataset to assess cross-dataset generalizability.
4. **SNP-trained model, ChIP-seq-tested**: A model was trained on the SNP dataset: 500 positives and 500 negatives) and tested on the 1000 ChIP-seq-derived sequences.

For negative sequence generation in all scenarios, we used the “getNullSeqs” function from the LS-GKM toolkit to obtain matched genomic background sequences, ensuring comparable GC content and sequence length. Each of the three experimental conditions was evaluated using three distinct methodologies: MinSeqChIP, MEME, and LS-GKM, allowing comparative analysis across different feature learning and classification frameworks. We ran MEME using the same parameters described in the section “MEME”. For LS-GKM, we used the default mode. For MinSeqChIP, the following settings were used: minimum monomer size set to 2, maximum monomer size set to 6, and a maximum allowed gap of 1 between monomers (due to the short and ungapped nature of the motifs). A Markov model of order 5 was applied in a non-gapped configuration. For PWM thresholding, we used the cutoff of 40 for the minimum number of sequences required to consider the library interpolated Markov model.

### Generating Flp-In 293 with stable NKLX2-5, GATA4, and TBX5 expression

Flp-In 293 cells (Thermo Fisher, #R75007) were modified to express TBX5 by inserting gene according to manufacturer instructions. In short, cells were seeded at 200,000 cell/mL in 2.5 mL of media (DMEM+10%FBS+0.1% Anti-anti) 2 days prior to transfection. 30 µL of Lipofectamine 3000® Reagent (ThermoFisher, #L300015) was diluted in 1mL Opti-MEM® (ThermoFisher, #31985062) and incubated for 5 minutes (37°C/5% CO2). A total of 2.5 µg of DNA (2.25 µg pOG44 + 0.25 µg of TBX5 gene insert) was diluted in 250 µL Opti-Mem. Both dilutions were combined to make a 1:1 ratio (250 μL Opti-MEM/Lipofectamine to 250 μL Opti-MEM/DNA) and incubated for 20 minutes (37°C/5% CO2). The old medium was removed from the cells and 500 µL of Opti-MEM/DNA/Lipofectamine mixture was added and incubated for 5 hours (37°C/5% CO2). Afterward, 1.5 mL of media was added to each well and incubated overnight. After reaching 80 % confluency, cells were switched to selection media containing hygromycin (50 µg/mL). Media was changed weekly to increase hygromycin until reaching 200 µg/mL. Once confluent, cells were transferred to a T75 plate until 80-90 % confluency for downstream experiments. Stocks were prepared at 3 million cells/mL in freezing media (HI-FBS (ThermoFisher, #A5670501) + 10% DMSO (Sigma Aldrich, #276855). Flp-In 293 cells already expressed NKX2-5 and GATA4 and were confirmed by RNA-seq and Western Blot.

### Massively Parallel Reporter Assay

For the assembly of the MPRA library, we followed the procedure described by Lu et al. with minor modifications (See MPRA method overview in Figure 4.2). ^47^ In brief, the MPRA library was cloned into a previously generated pGL4.23ΔxbaΔluc empty vector from pGL4.23 [luc2/minP] (Promega, # E8411). First, 20 bps barcodes were added to the synthesized oligos through 24X PCR with 50 μL system, each containing 1.86 ng oligo, 25 μL NEBNext® Ultra™ II Q5® Master Mix (NEB, #M0544S), 1 μM MPRA_v3_F, and 1 µM MPRA_v3_201_R. PCR was performed under the following conditions: 98 °C for 2 min, 12 cycles of (98 °C for 10 s, 60 °C for 15 s, 72 °C for 45 s), 72 °C for 5 min. The amplified product was purified with 1.8X Ampure XP SPRI beads (Beckman Coulter, A63881) and cloned into SfiI-digested pGL4.23ΔxbaΔluc by Gibson assembly (1:2 molar ratio of vector:insert) at 50 °C for 1 h. The assembled backbone library was purified and then transformed into 10-β Escherichia coli (*E. coli*; NEB, #C3019H) through electroporation (2 kV, 200 ohm, 25 μF) according to manufacturer instructions. Electroporated *E. coli* was expanded in 200 mL of LB Broth supplemented with 100 μg/mL of carbenicillin at 37°C for 12 to 16 h. Plasmid was then extracted using the QIAGEN Plasmid Maxi Kit (Qiagen, #12162).

To associate barcodes with oligo sequences, 200ng of the plasmid was amplified using NEBNext Q5® High-Fidelity 2X Master Mix (NEB, #M0492L), with primers TruSeq_Universal_Adapter and MPRA_v3_TruSeq_Amp2Sa_F (Table S7) using the following conditions: 95°C for 20s, 6 cycles (95°C for 20s, 62°C for 15s, and 72°C for 30s), and 72°C for 2 minutes. The PCR product was purified using SPRI beads at 0.5X and then 1X. Next we performed additional 5 cycles of PCR to attached custom Illumina P7 indices purified using SPRI beads at 0.5X and then 1X. Samples were sequenced on a NovaSeq X Plus (2 × 150 bp) at Cincinnati Children’s Hospital sequencing core to achieve a 30X read coverage. Identification of which barcodes were associated with which oligos was then conducted with the MPRAmatch pipeline (https://github.com/tewhey-lab/MPRAmatch).

A miniP + eGFP fragment was amplified from pGL4.23[eGFP/miniP] through 20X PCR with 50 μL system, each containing 1 ng plasmid, 25 μL NEBNext® Ultra™II Q5® Master Mix, 0.5 μM 200-MPRA_v3_GFP_Fusion_v2_F, and 201-MPRA_v3_GFP_Fusion_v2_R. PCR was performed under the following conditions: 98 °C for 2 min, 20 cycles of (98 °C for 10 s, 60 °C for 15 s, 72 °C for 45s), 72 °C for 5 min. The amplified product was purified with 1.8X SPRI and then inserted into AsiSI digested pGL4:23:ΔxbaΔluc backbone library through Gibson assembly at 50 °C for 1.5 h and purified at 1.5X SPRI. The resulting library was re-digested by RecBCD (NEB, #M0345S) and AsiSI (NEB, #R063L), purified at 1.5X SPRI, and then transformed into *E. coli* through electroporation (2 kV, 200 ohm, 25 μF). Transformed *E. coli* were cultured in 5 L of LB Broth supplemented with 100 μg/mL of carbenicillin at 37 °C for 12–16 h. The plasmid was then extracted using the QIAGEN Endo-free Plasmid Giga Kit (Qiagen, #12191).

### Transfection of MPRA Library

Flp-In 293 cells (ThermoFisher, #R75007) were cultured in 30 mL Dulbecco’s modified Eagle’s medium (DMEM) (ThermoFisher, #10564) containing 10% fetal bovine serum (FBS) (ThermoFisher, #A3160401) and 1% penicillin-streptomycin (Pen-Strep) (ThermoFisher, #15140122). Five total replicates were transfected. For each replicate, cells were plated in two 15-cm plates and grown to a density of ∼80 to 90% (∼20 to 40 million cells per plate). Cells were then incubated with 80 μL of Lipofectamine 3000 (ThermoFisher, L3000015) and 20 μg of DNA for 24 hours. Then, transfected cells were split 1:3 into new 15-cm plates, keeping all transfected cells. After an additional 24 hours (48 hours after transfection), cells were pelleted by centrifugation, washed once with phosphate-buffered saline (PBS), flash-frozen using liquid nitrogen, and then stored at –80°C.

### MPRA RNA Sample Processing

RNA processing was performed as previously described. ^168^ In short, total RNA was extracted from the cell pellets with the Qiagen Maxi RNeasy kit (Qiagen, #75162) with on-column DNase digest according to manufacturer’s instructions. A DNase reaction was further performed to remove remaining MPRA library vectors using Turbo DNAse kit (ThermoFisher, #AM2238). The reaction was stopped with 0.1% SDS (Sigma Aldrich, #71736) and 0.05M EDTA (Sigma Aldrich, #03690). The GFP-transcripts in total RNA were then captured through a hybridization reaction with streptavidin beads (ThermoFisher, #65001) and three GFP-targeted biotinylated oligos (Table S7). RNA was then cleaned up with RNA SPRI (Beckman Coulter, #A63987) and converted to cDNA using a Superscript III (ThermoFisher, 18080044) reaction with primer MPRA_v3_Amp2Sc_R (Table 4.1). The relative cDNA abundance was estimated through quantitative PCR along with serial dilutions of plasmid library serving as a standard curve (see Table S7 for primer sequences). The PCR conditions were: 98°C for 30s, 40x of (95°C for 20s, 65°C for 20s, and 72°C for 30s), and 72°C for 2 minutes. To minimize amplification bias, the Ct number reflecting the point at which the amplification just began to take off, subtracted by 1, was used to set up the first PCR for sequencing preparation. cDNA and plasmids were normalized to approximately the same concentration and cycled for 10 cycles using NEBNext Q5® High-Fidelity 2X Master Mix (NEB, #M0492L) and primers MPRA_v3_Illumina_GFP_F and TruSeq_Universal_Adapter (Table 4.1). The product was cleaned up with RNA SPRI at 1X, eluted in 30μL, 20μL of which was then subjected to another round of 6 cycles to attach custom p7 and p5 Illumina adapters with unique sample indices. Samples were sequenced on a NextSeq 2000 platform using the P3 100 cycle kit, with an average of around 100M reads per sample.

### MPRA Data Analysis

Oligo counts were obtained via the MPRAcount pipeline (https://github.com/tewhey-lab/MPRA_oligo_barcode_pipeline). Oligos with at least 10 barcodes were retained for analysis and oligo counts were normalized for sequencing depth with the DESeq2 median of ratios method. DESeq2 was then used to estimate the fold change between plasmid DNA and cDNA with Wald’s test and p-values were corrected for multiple hypothesis testing by Bonferroni’s method. Significance threshold was determined at adjusted p-value less than 0.01 in either the reference or alternate allele in order to call a sequence as having a regulatory effect on expression. For identification of expression-modulating variants, only variants originating from sequences determined to have a regulatory effect were considered. Allelic skew was calculated by comparing the log ratios of the reference and alternative alleles using Wald’s test. All skew p-values were adjusted with the Benjamini-Hochberg procedure and determined to be significant at 5% false discovery rate. The scripts for estimating variant activity and allelic skew are available on https://github.com/tewhey-lab/MPRAmodel. Scripts for plotting and visualization of MPRA analysis are available on https://github.com/WeirauchLab/mpraprofiler.

### HOMER motif enrichment analysis

HOMER (v5.1) ^90^ was performed on 50 bp reference and alternate fasta file centered on the CVD-associated SNP using the findMotif.pl function. The background file was generated using the homer2 background function. Shared motifs between a reference allele and its corresponding alternate allele(s) were removed from the analysis. This ensured that only motifs that occur in only the reference or alternate allele were used for downstream analysis. Finally, TF and family counts were generated by counting unique instances of SNP-TF/family pairs for either the reference or alternate allele. Additionally, we used HOMER to calculate the enrichment of each motif in the sequence of emAlleles compared to the sequences of non-emAlleles.

### Functional genomics dataset enrichment analysis with RELI

We used the Regulatory Element Locus Intersection (RELI) ^74^ algorithm to identify genomic features (TF binding events, histone marks, etc.) that coincide with emAlleles as previously described. ^47^ In short, RELI systematically intersects the coordinates of emAlleles with a large collection of ChIP-seq datasets and counts the number of times the input regions overlap with peaks in the dataset by at least one base. A p-value describing the significance of this overlap is estimated by comparing the input set with a negative set (variants with no CRE activity; 50% expression fold change). We performed the RELI analysis for all variants categorized as emAlleles, and for the individual subset that overlapped with cardiac CREs.

## Key Resources Table

Key Resources Table:
| REAGENT or RESOURCE | SOURCE | IDENTIFIER |
| --- | --- | --- |
| Antibodies |  |  |
| α-NKX2-5 | Cell Signaling | 8792 |
| α-GATA4 | Cell Signaling | 36966 |
| α-FLAG | Sigma Aldrich | F1804-50UG |
| Bacterial and virus strains |  |  |
| 10-β E. Coli | NEB | C3019H |
| Chemicals, peptides, and recombinant proteins |  |  |
| KAPA HiFi HotStart ReadyMix | Roche Sequencing Store | KK2602<br>07958935001 |
| Sodium Chloride | Fisher Scientific | S671-3 |
| UltraPure® 1M Tris-HCl, pH 8.0 | Invitrogen | 15568025 |
| Glycerol | Sigma Aldrich | G5516 |
| pdl-dC | Sigma Aldrich | P4929 |
| BSA | Fisher Scientific | 23210 |
| Tween-20 | Sigma Aldrich | P9416 |
| DTT | Sigma Aldrich | D0632 |
| HEPES | VWR | 7365-45-9 |
| Zinc Acetate | Sigma Aldrich | 383058 |
| 10X TBE(Tris/Boric Acid/ EDTA) | Biorad | 1610770 |
| Buffer EB | Qiagen | 19086 |
| Dynabeads M-280 Streptavidin | ThermoFisher | 11205D |
| DMEM | ThermoFisher | 10564 |
| FBS | ThermoFisher | A3160401 |
| Opti-MEM | ThermoFisher | 31985062 |
| Lipofectamine 3000 Transfection Reagent | ThermoFisher | L300015 |
| Hygromycin B | Sigma Aldrich | H3274 |
| HI-FBS | ThermoFisher | A5670501 |
| DMSO | Sigma Aldrich | 276855 |
| NEBNext® Ultra™ II Q5® Master Mix | NEB | M0544S |
| Gibson Assembly Master Mix | NEB | E2611L |
| XbaI | NEB | R0145S |
| NcoI | NEB | R0193S |
| RecBCD | NEB | M0345S |
| AsiSI | NEB | R063L |
| NEBNext Q5® High-Fidelity 2X Master Mix | NEB | M0492L |
| Penicillin-Streptomycin | ThermoFisher | 15140122 |
| SDS | Sigma Aldrich | 71736 |
| EDTA | Sigma Aldrich | 03690 |
| Streptavidin Beads | ThermoFisher | 65001 |
| RNA SPRI | Beckman Coulter | A63987 |
| Superscript III | ThermoFisher | 18080044 |
| Critical commercial assays |  |  |
| Q5 Site-directed Mutagenesis Kit | NEB | E0554S |
| Qiagen Plasmid Maxi Kit | Qiagen | 12162 |
| Qiagen Endo-free Plasmid Giga Kit | Qiagen | 12191 |
| Qiagen Maxi RNeasy | Qiagen | 75162 |
| Turbo DNase Kit | ThermoFisher | AM2238 |
| Deposited data |  |  |
| CHD variant dataset from the GWAS catalog | Ref 49-54 | N/A |
| ChIP-seq Dataset NKX2-5 | Gonzalez-Terran et al<br>Cell 2022 | SRX9284027 |
| ChIP-seq Dataset GATA4 | Gonzalez-Terran et al<br>Cell 2022 | SRX9284038 |
| ChIP-seq Dataset TBX5 | Ang, Y.S. et al Cell<br>2016 | SRX2023721 |
| CHD variants MPRA dataset | GEO LINK |  |
| CHD variants SNP Bind-n-Seq dataset | GEO LINK |  |
| Experimental models: Cell lines |  |  |
| Flp-In 293 | ThermoFisher | R75007 |
| Oligonucleotides |  |  |
| MPRA_v3_F | IDT | N/A |
| MPRA_v3_201_R | IDT | N/A |
| GFP_seq_MS2_P65-HSF1_GFP_FWD | IDT | N/A |
| GFP_seq_MS2_P65-HSF1_GFP_REV | IDT | N/A |
| 200-MPRA_v3_GFP_Fusion_v2_F | IDT | N/A |
| 200-MPRA_v3_GFP_Fusion_v2_R | IDT | N/A |
| TruSeq_Universal_Adapter | IDT | N/A |
| MPRAv3_a2sa | IDT | N/A |
| MPRA_v3_Amp2Sc_R | IDT | N/A |
| MPRA_v3_Illumina_GFP_F | IDT | N/A |
| Recombinant DNA |  |  |
| Oligopool of CHD-associated variants | Custom Array | N/A |
| MPRA Oligopool | Twist Bioscience | N/A |
| pET-NKX2.5-6XHisTag | VectorBuilder, Inc. | N/A |
| pET-TBX5-6XHisTag | VectorBuilder, Inc. | N/A |
| pET-28a(+)-6XHisTag-GATA4 DBD | Twist Bioscience | N/A |
| pOG44 | ThermoFisher | V600520 |
| pGL4.23 [Luc2/minP - firefly luciferase] | Promega | E8411 |
| MS2-P65-HSF1_GFP | Addgene | 61423 |
| Software and algorithms |  |  |
| TopLD Web Tool | Yun Li Lab<br>http://topld.genetics.unc.edu/ | N/A |
| Plink (v1.90b) |  | N/A |
| UCSC Genome Browser |  | N/A |
| Scipy.optimize (Python) |  | N/A |
| MEME Motif Discovery | https://meme-suite.org/meme/ | N/A |
| MinSeqChIP |  | N/A |
| LS-GKM |  | N/A |
| MPRAMatch pipeline | Tewhey Lab | N/A |
| DESeq2 |  | N/A |
| MPRAcount pipeline |  | N/A |
| Homer(v5.1) | Ref82 | N/A |
| Regulatory Element Locus Intersection (RELI) | Ref66 | N/A |
| Other |  |  |
| Azure Zaphire Biomolecular Imager | Azure Biosystems | N/A |
| Thermoshaker | BIOGRANT | N/A |

## Supplementary Figures

**Supplementary Figure 1:**
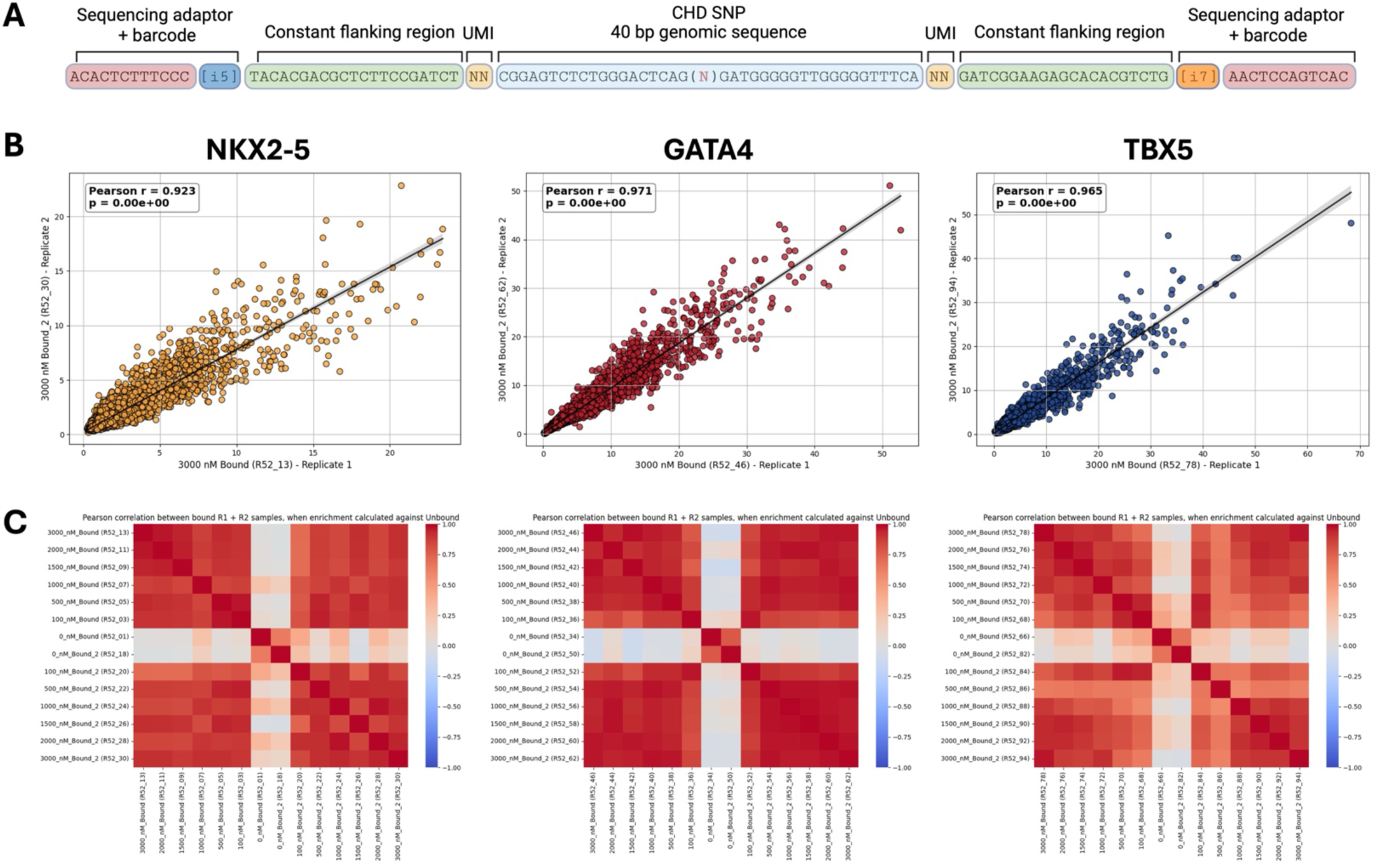
SNP Bind-n-Seq library anatomy and replicate correlation. **A)** SNP Bind-n-Seq library sequence features, structures, and constant regions. **B)** Experimental correlation between replicates of NKX2-5 (left), GATA4 (middle), and TBX5 (right) at 3,000 nM. **C)** Correlation Matrix of Replicate Enrichment Values Relative to Unbound Conditions. Heatmap of the correlation matrix of NKX2-5 (left), GATA4 (middle), and TBX5 (right) enrichment across concentrations, relative to the corresponding Unbound condition in both replicates. Each cell represents the Pearson correlation coefficient between two concentrations, indicating the similarity in enrichment patterns.

**Supplementary Figure 2:**
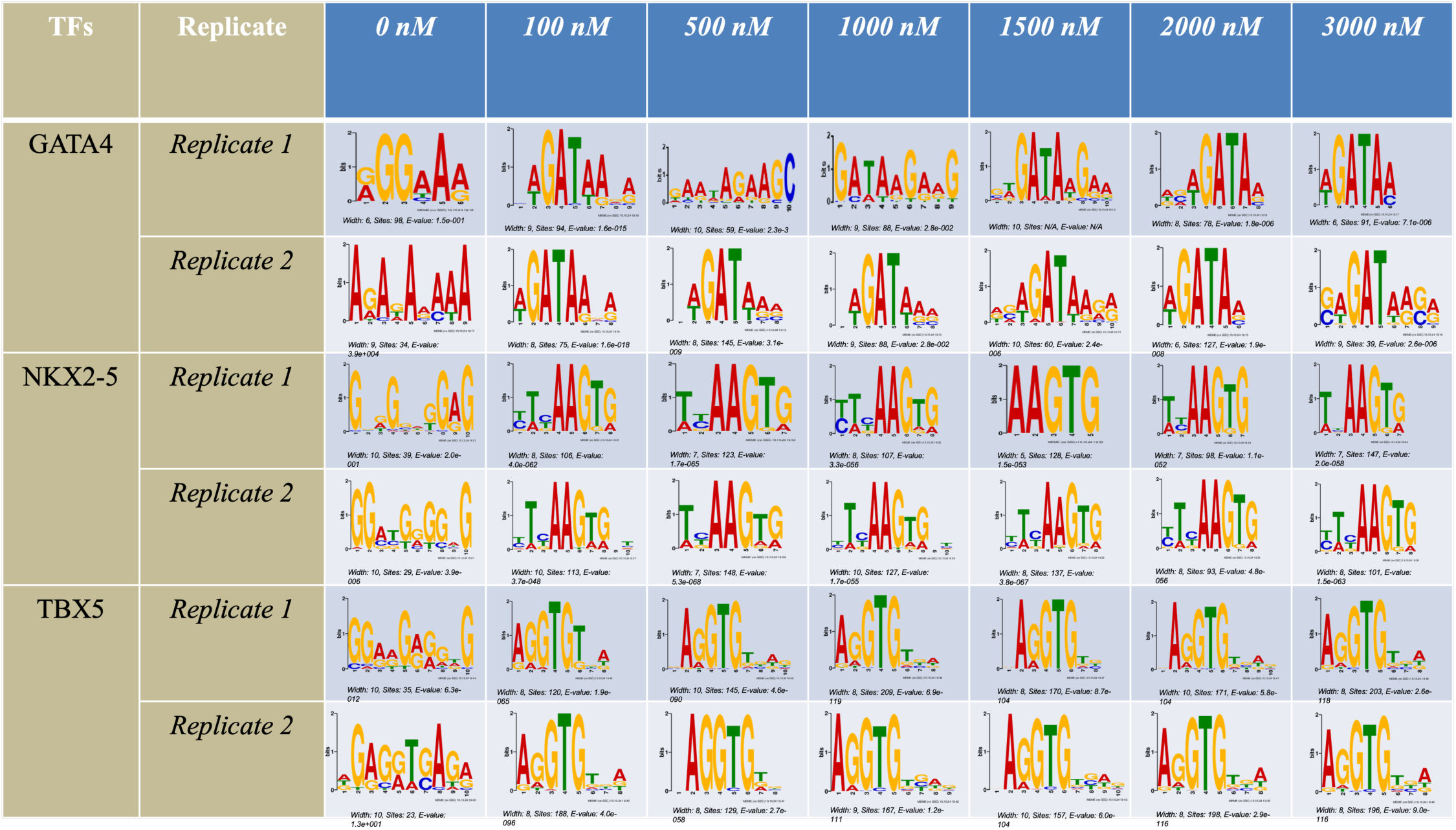
TF motif enrichment across SNP Bind-n-Seq replicates and concentration points. Motifs were generated using MEME with the top 500 sequences with the highest KA values.

**Supplementary Figure 3:**
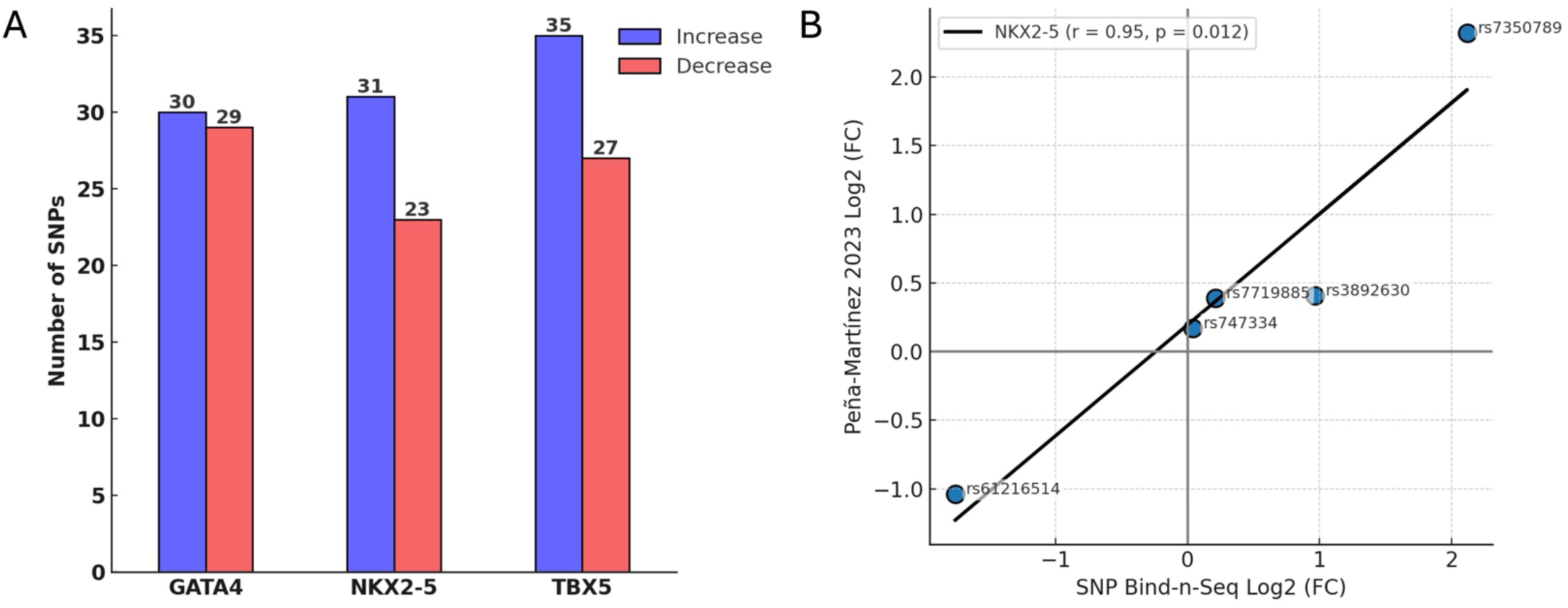
Variants with differential allelic binding identified through SNP Bind-n-Seq. **A)** Number of variants with allelic binding for NKX2-5, GATA4, and TBX5. **B)** Fold change correlations of previously described variants for NKX2-5 binding affinity.

**Supplementary Figure 4:**
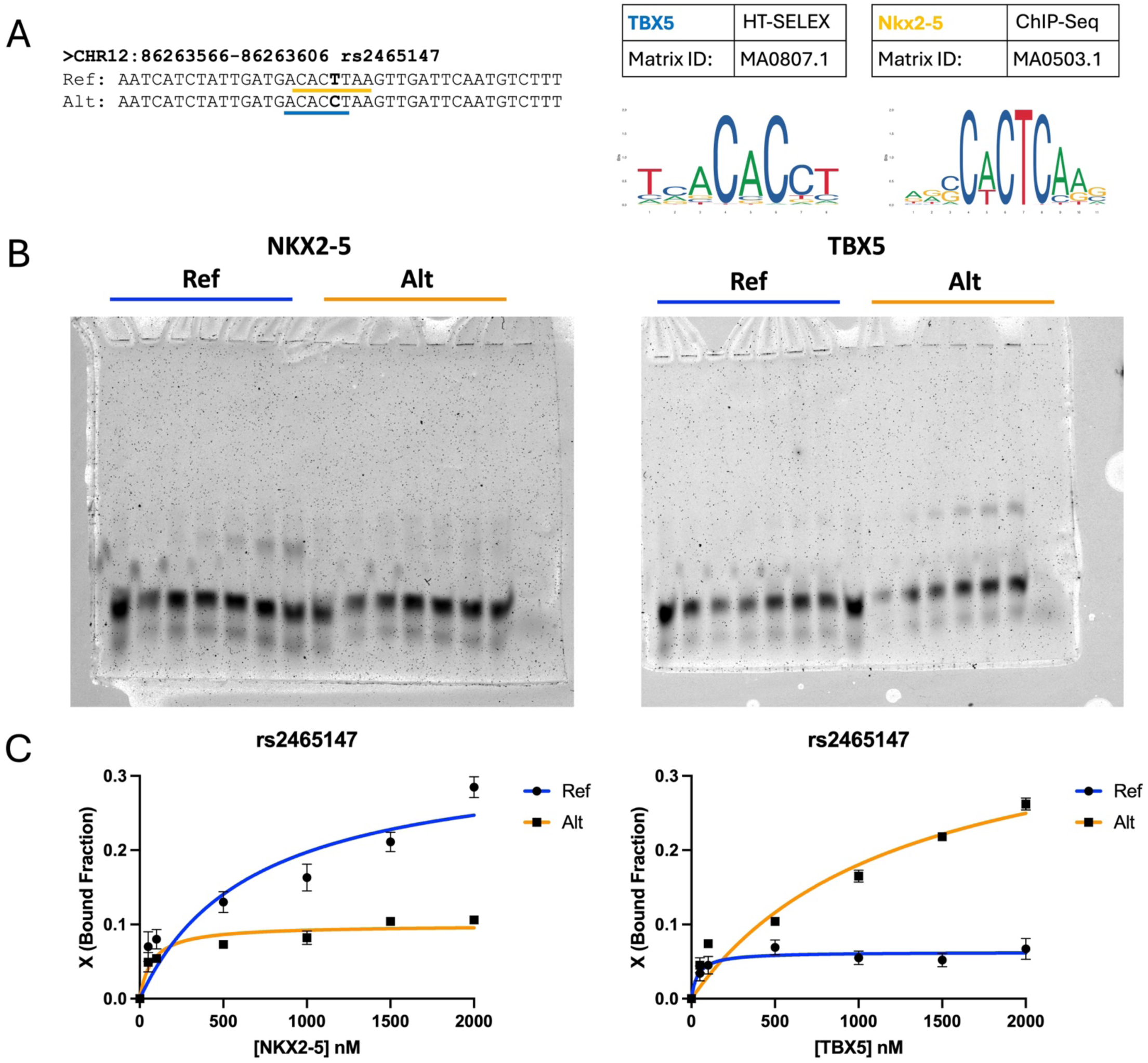
In vitro validation of SNP Bin-n-Seq variant rs2465147. **A)** Reference and alternate sequences. TF binding motifs are highlighted in yellow for NKX2-5 and blue for TBX5. JASPAR motifs from NKX2-5 and TBX5 are displayed on the right. **B)** Electrophoretic mobility shift assay (EMSA) of rs2465147 for NXK2-5 (left) and TBX5 (right). **C)** Binding curves generated from EMSA of rs2465147 in triplicate.

**Supplementary Figure 5:**
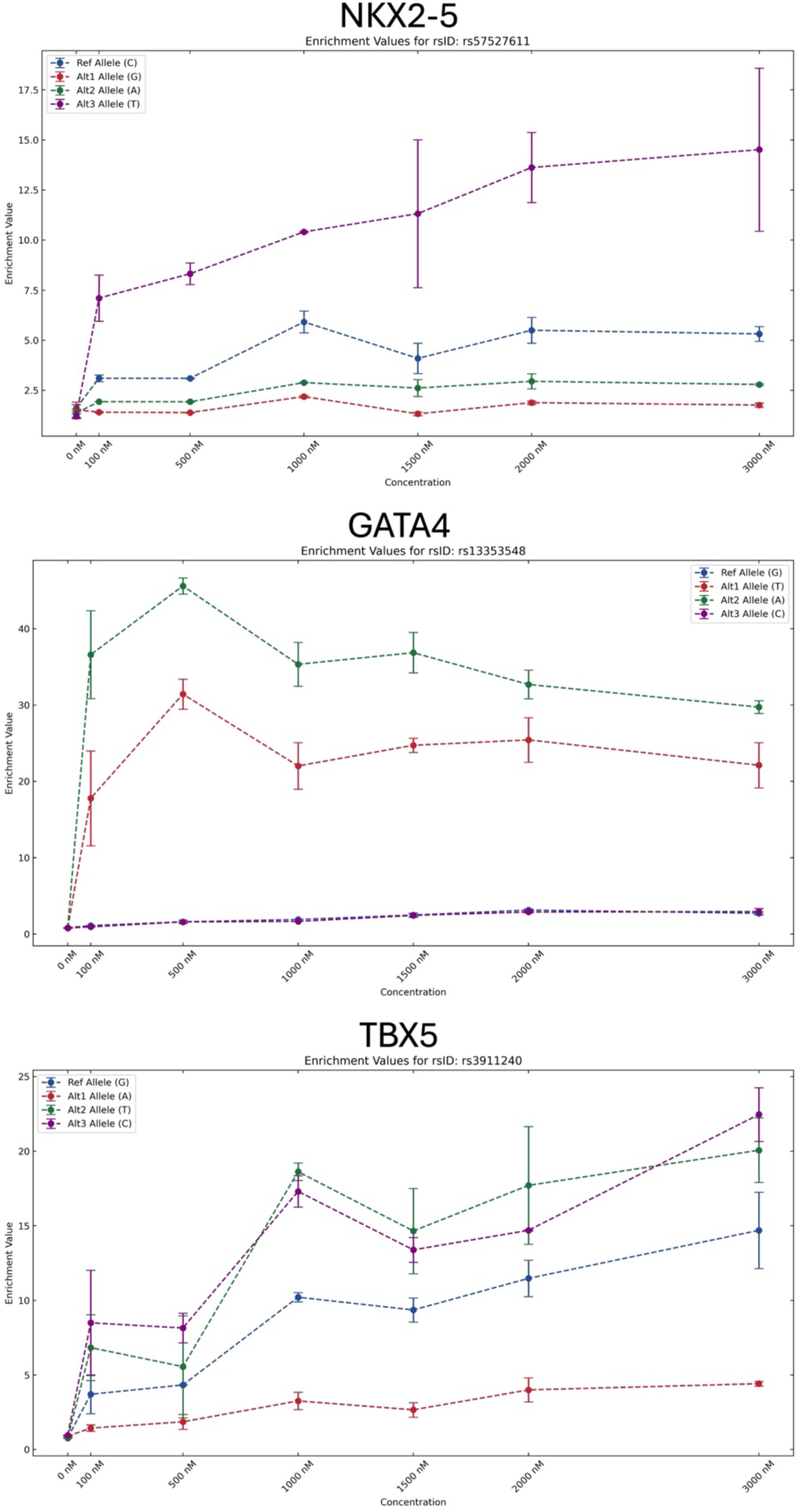
Allelic enrichment curve of non-CHD-risk alternate alleles. Variants rs57527611, rs13353548, and rs3911240 are plotted for NKX2-5 (top), GATA4 (middle), and TBX5 (bottom) binding, respectively. Reference alleles (Ref) are represented in blue, and tag-SNP alleles from the GWAS catalog (Alt 2) are represented in red. Permutated alleles (alternate non-risk; Alt 2 and Alt 3) are represented in green and purple, respectively.

**Supplementary Figure 6:**
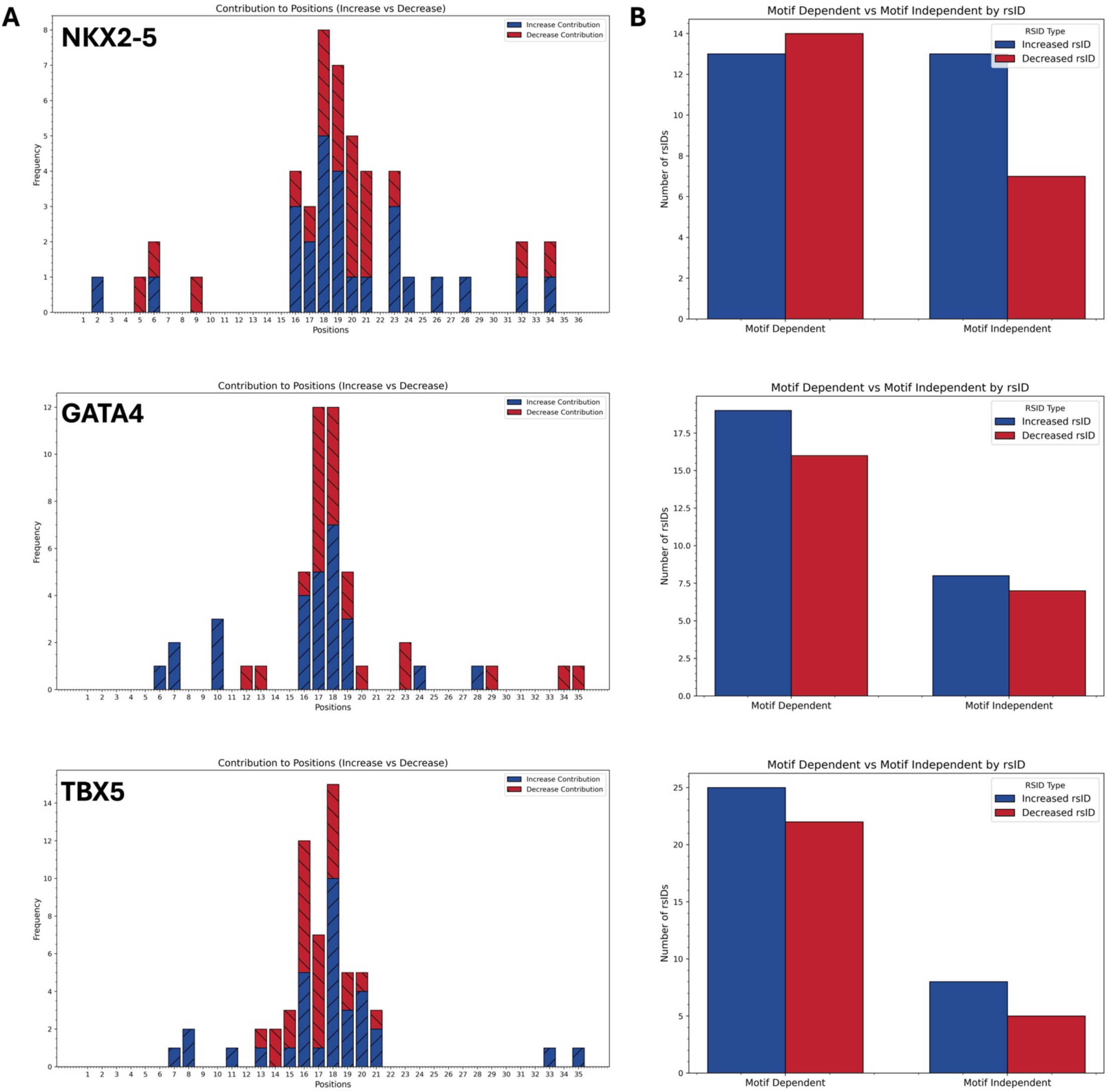
Motif disruption analysis of variants with allele-specific binding. **A)** Binding motifs distribution of variants with allele-specific binding. X-axis represents the 40 bp window centered on the variant. **B)** Count of variants that directly create or disrupt TF binding motifs (motif dependent), adjacent to TF motif (motif independent). Variants creating binding motifs are represented in blue, whereas disrupting variants are represented in red.

**Supplementary Figure 7:**
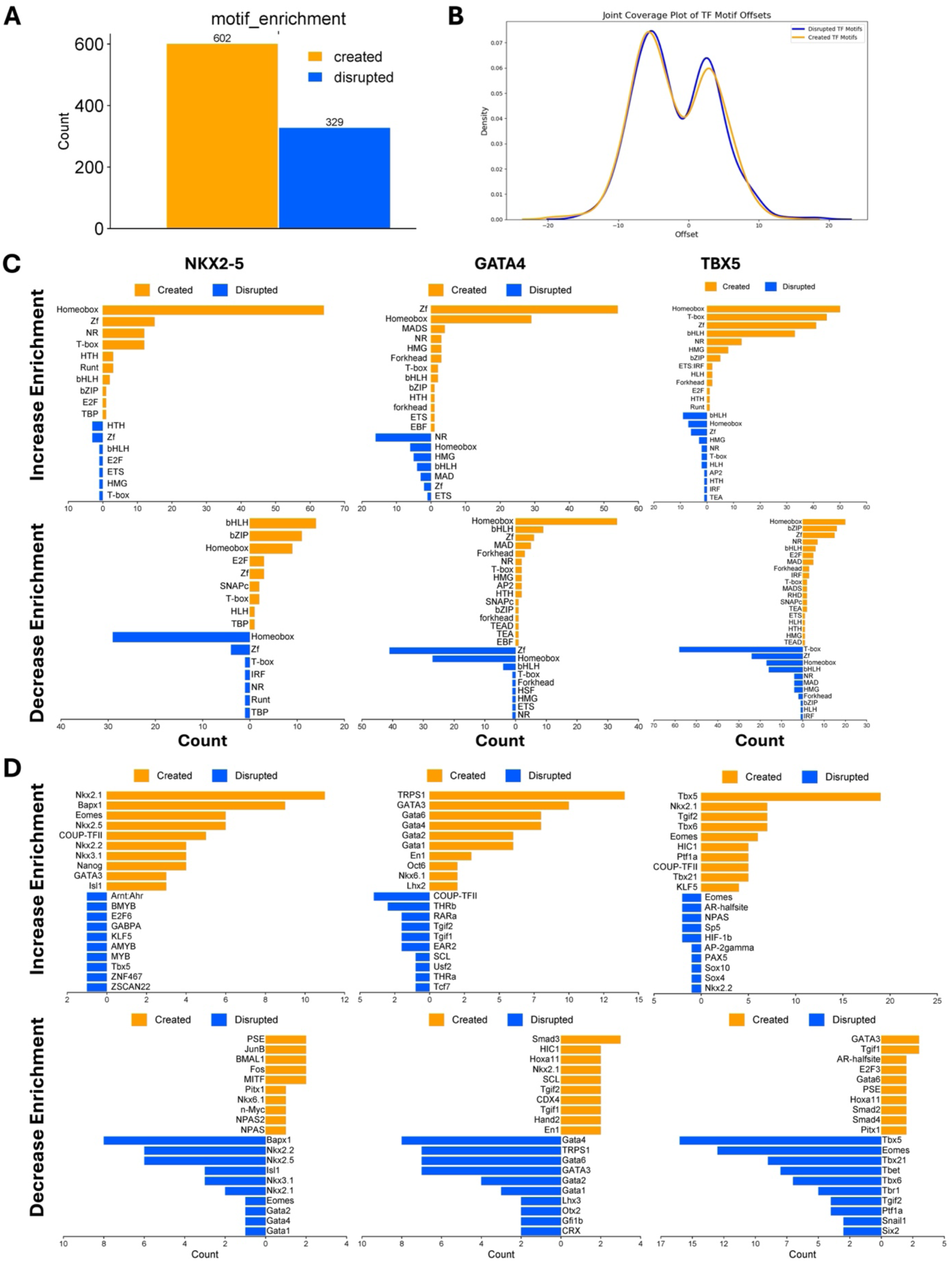
Homer motif enrichment analysis of variants with allele-specific binding. A) Number of variants that created (orange) or disrupted (blue) TF binding motifs. B) Frequency of created and disrupted TF binding motifs relative to variant position (X = 0). C-D) Number of motif created or disrupted for C) TF families and D) specific TFs. Motif enrichment analysis are displayed separately for variants that increased or decreased binding affinity for NXK2-5, GATA4, and TBX5.

**Supplementary Figure 8:**
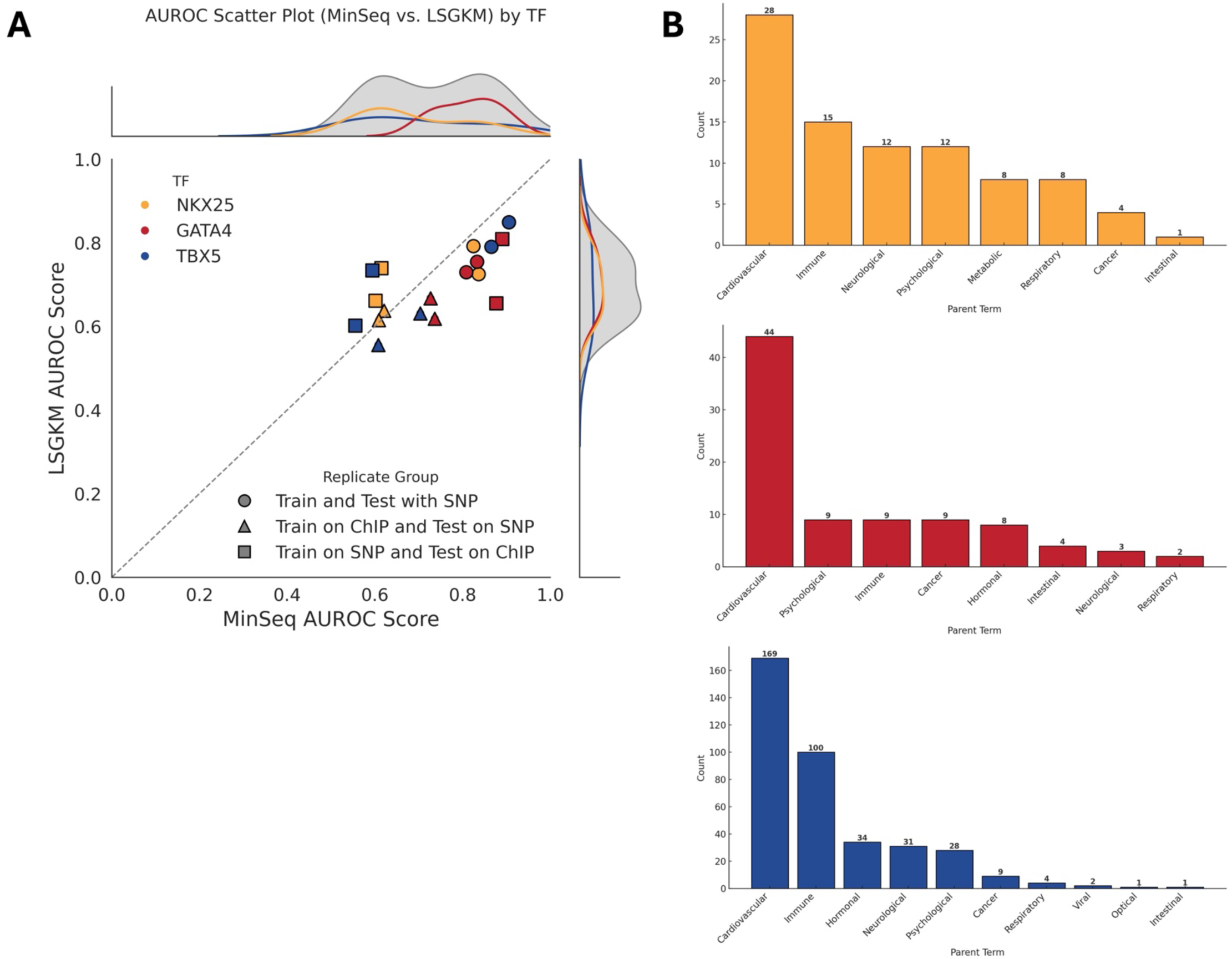
Training predictive models with SNP Bind-n-Seq experimental data. **A)** Scatter plot comparing MinSeqChIP and LSGKM classifier performance for three transcription factors (NKX2-5, GATA4, TBX5), with AUROC scores shown on the x-axis (MinSeqChIP) and y-axis (LSGKM). Marker shapes indicate evaluation strategies: circles represent training and testing on SNPs, triangles indicate training on ChIP-seq and testing on SNPs, and squares indicate training on SNPs and testing on ChIP-seq. For each condition, two points are plotted because the same positive set was evaluated against two different negative sets: LSGKM-generated negatives and genomic negatives located 6,000 bp away from positives. In SNP→SNP evaluation, a 60–40 split is used for both classifiers, with 60% of the data (300 positives and 300 negatives) used for training and 40% (200 positives and 200 negatives) used for testing. In ChIP→SNP evaluation, training is performed on the top 1,000 ChIP-seq peaks and testing on 500 SNP positives (controls excluded). In SNP→ChIP evaluation, training is on 500 SNP positives (controls excluded) and testing on 1,000 ChIP-seq peaks. The dashed diagonal line indicates equal classifier performance, while marginal kernel density estimates summarize the overall distribution of AUROC scores. Points above the diagonal indicate superior LSGKM performance, whereas points below indicate better MinSeqChIP performance. **B)** Number of variants predicted to alter TF binding per disease parent term from the GWAS catalog. NKX2-5 is represented in yellow, GATA4 in red, and TBX5 in blue.

**Supplementary Figure 9:**
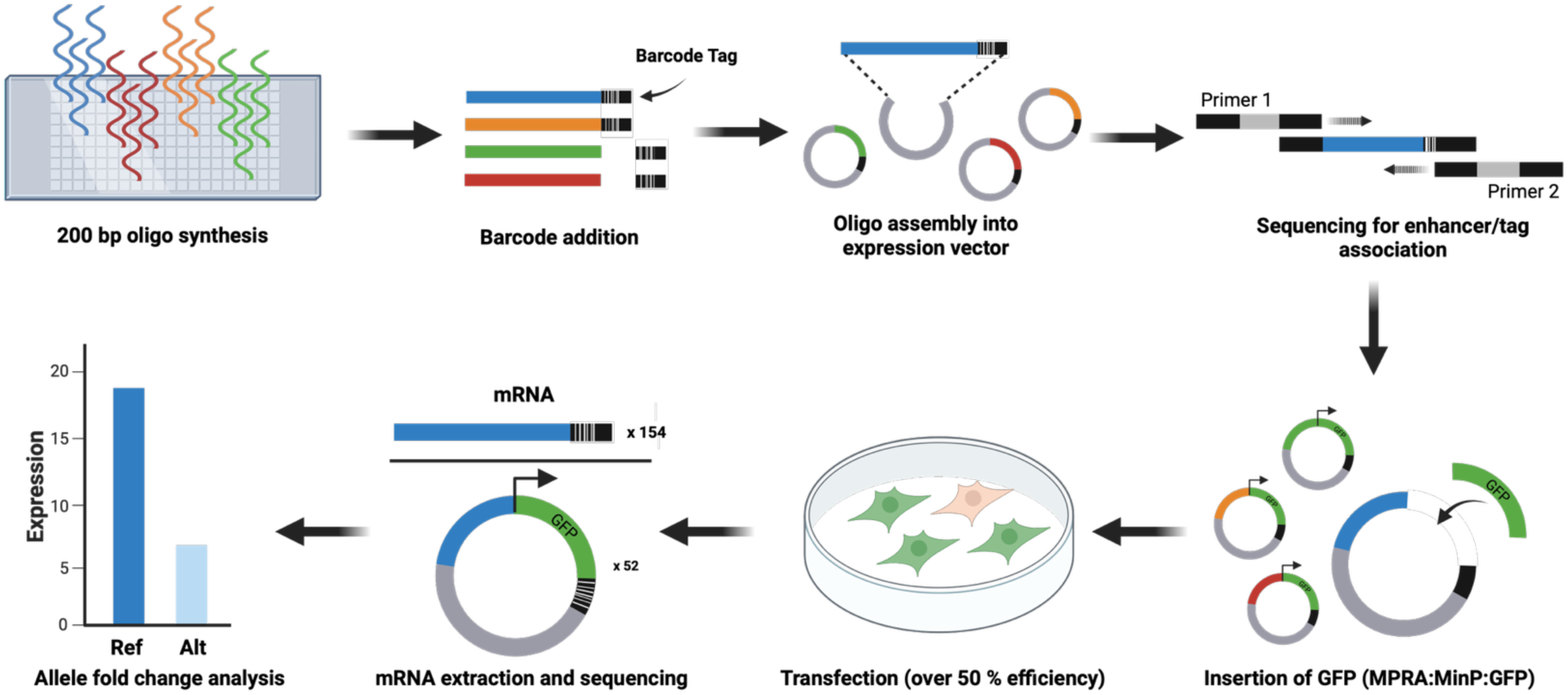
MPRA workflow.

**Supplementary Figure 10:**
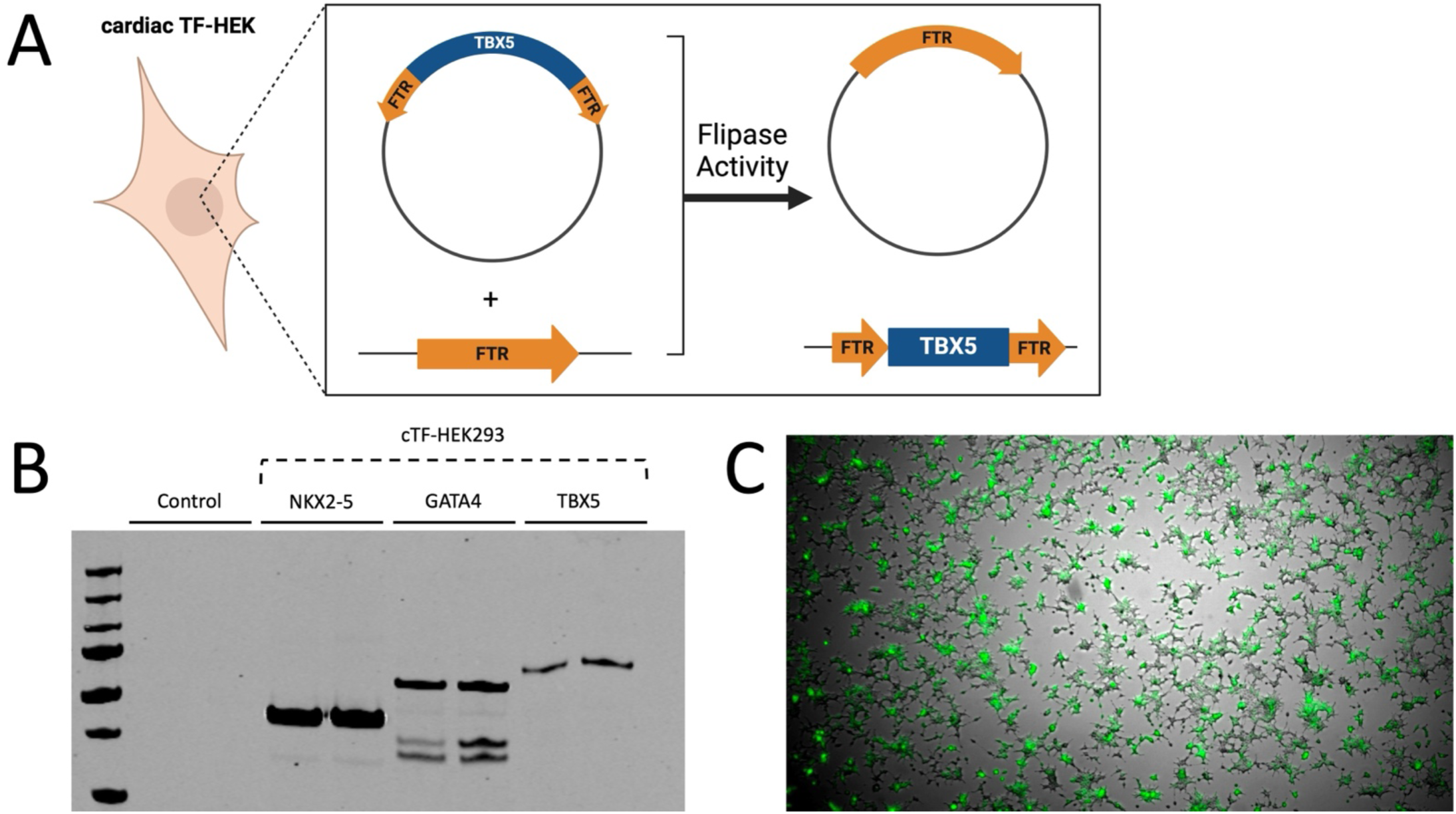
Generating a cardiac TF stably-expressing HEK293 cell line. **A)** Diagram of the HEK293 FlpIn system to integrate the cardiac TF gene into the genomic landing pad. **B)** Confirmation of NXK2-5, GATA4, and TBX5 expression in HEK FlpIn cell line through Western Blot. **C)** GFP library mock transfection of HEK FlpIn.

**Supplementary Figure 11:**
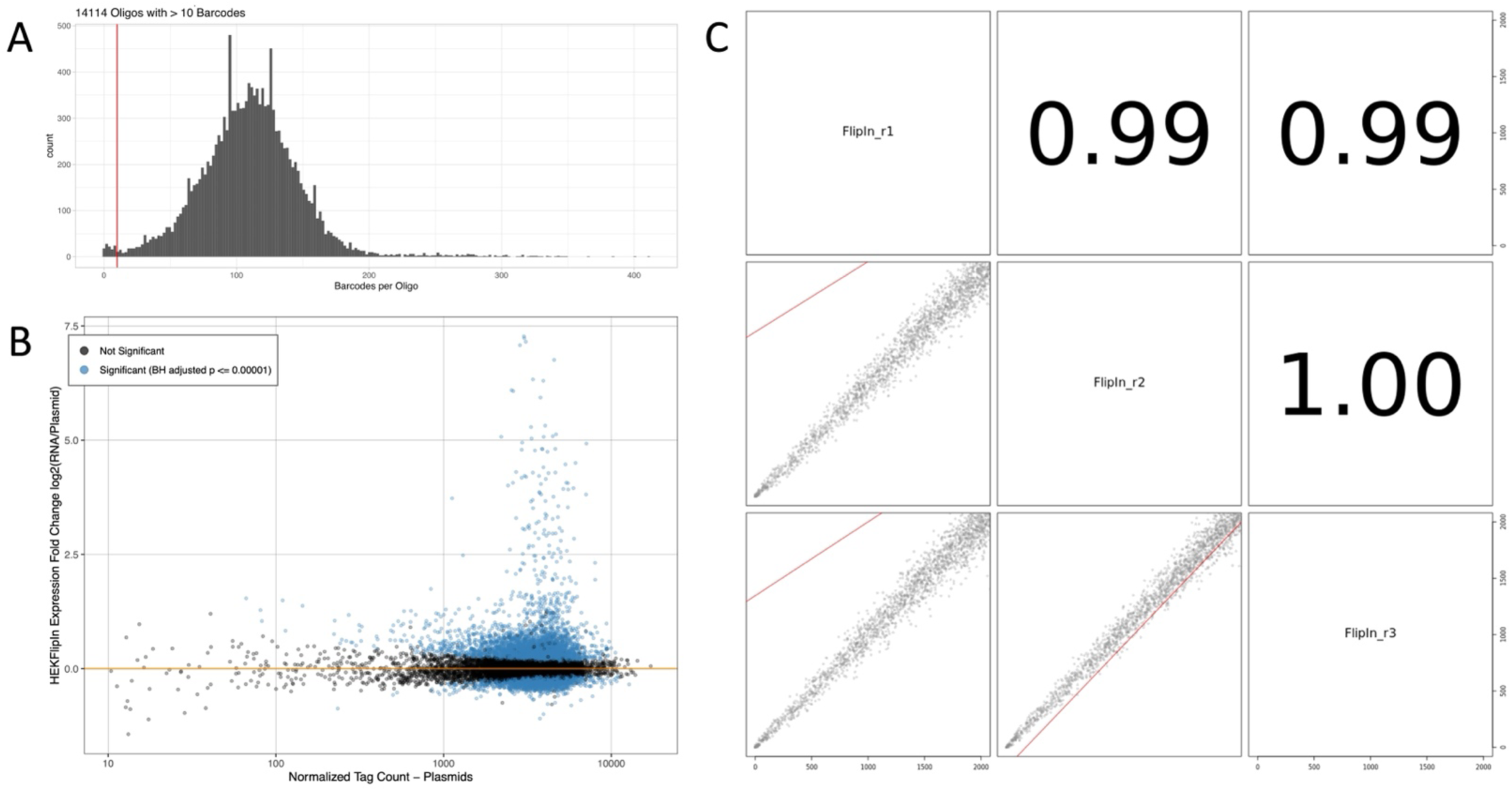
MPRA quality and reproducibility analysis. A) Number of variants with >10 unique barcodes per oligo. B) Fold change of oligos compared to barcode tag counts. C) Correlation between biological triplicates.

**Supplementary Figure 12:**
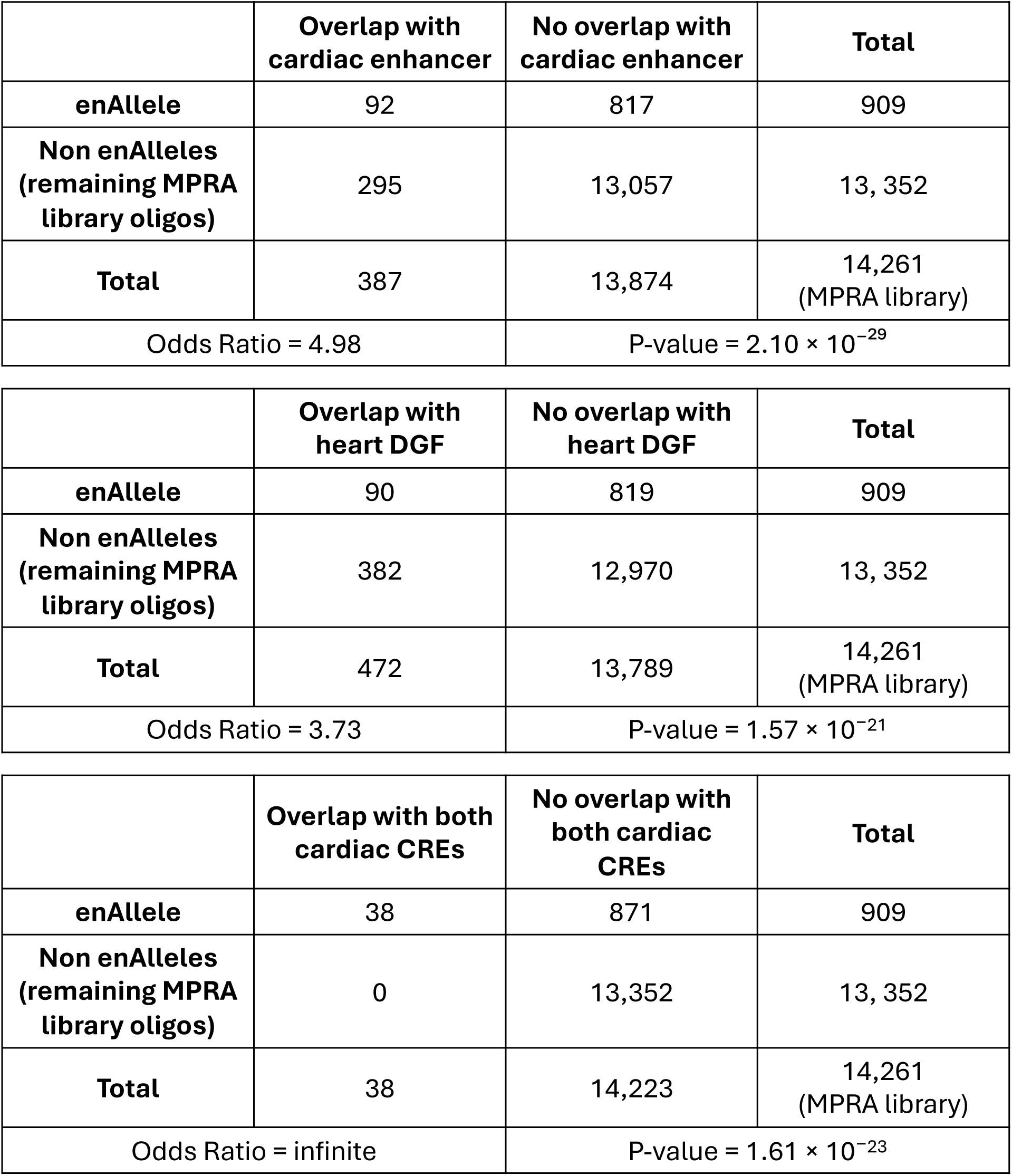
Contingency tables to determine the significance of overlap between enhancer alleles (enAlleles) with cardiac regulatory elements.

**Supplementary Figure 13:**
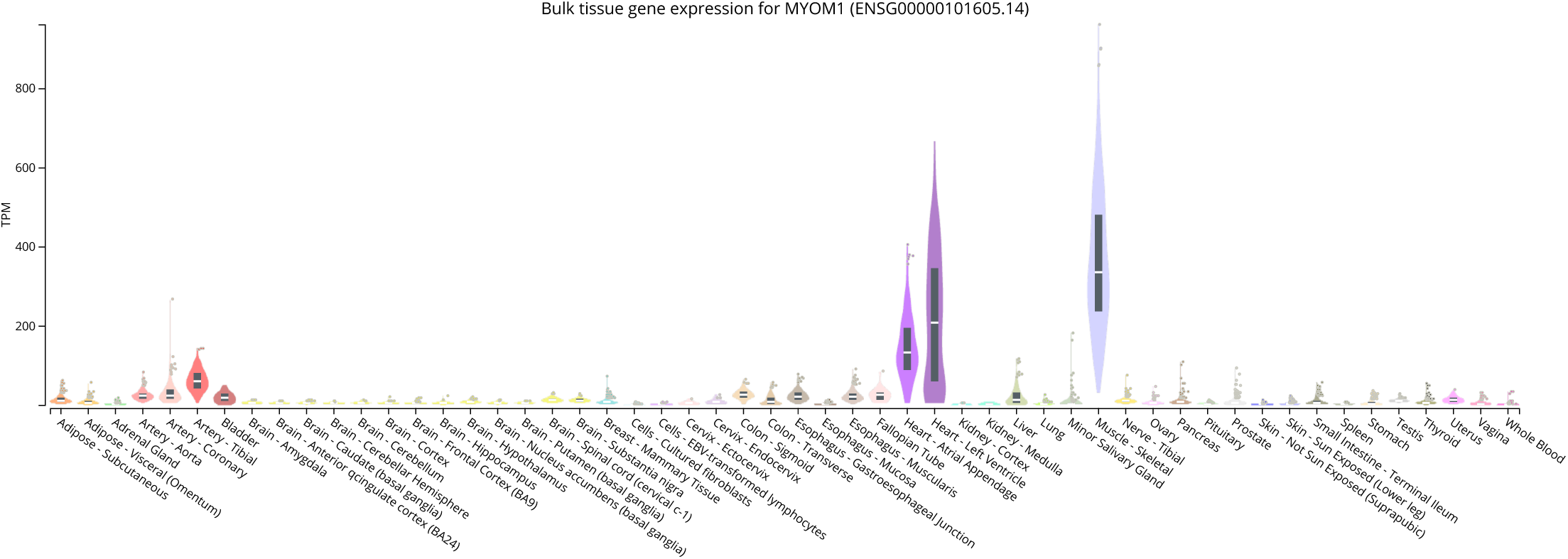
MYOM1 expression profiles in multiple tissues. Cardiac tissues (the heart atrial appendage and left ventricle) are displayed in purple.

**Supplementary Figure 14:**
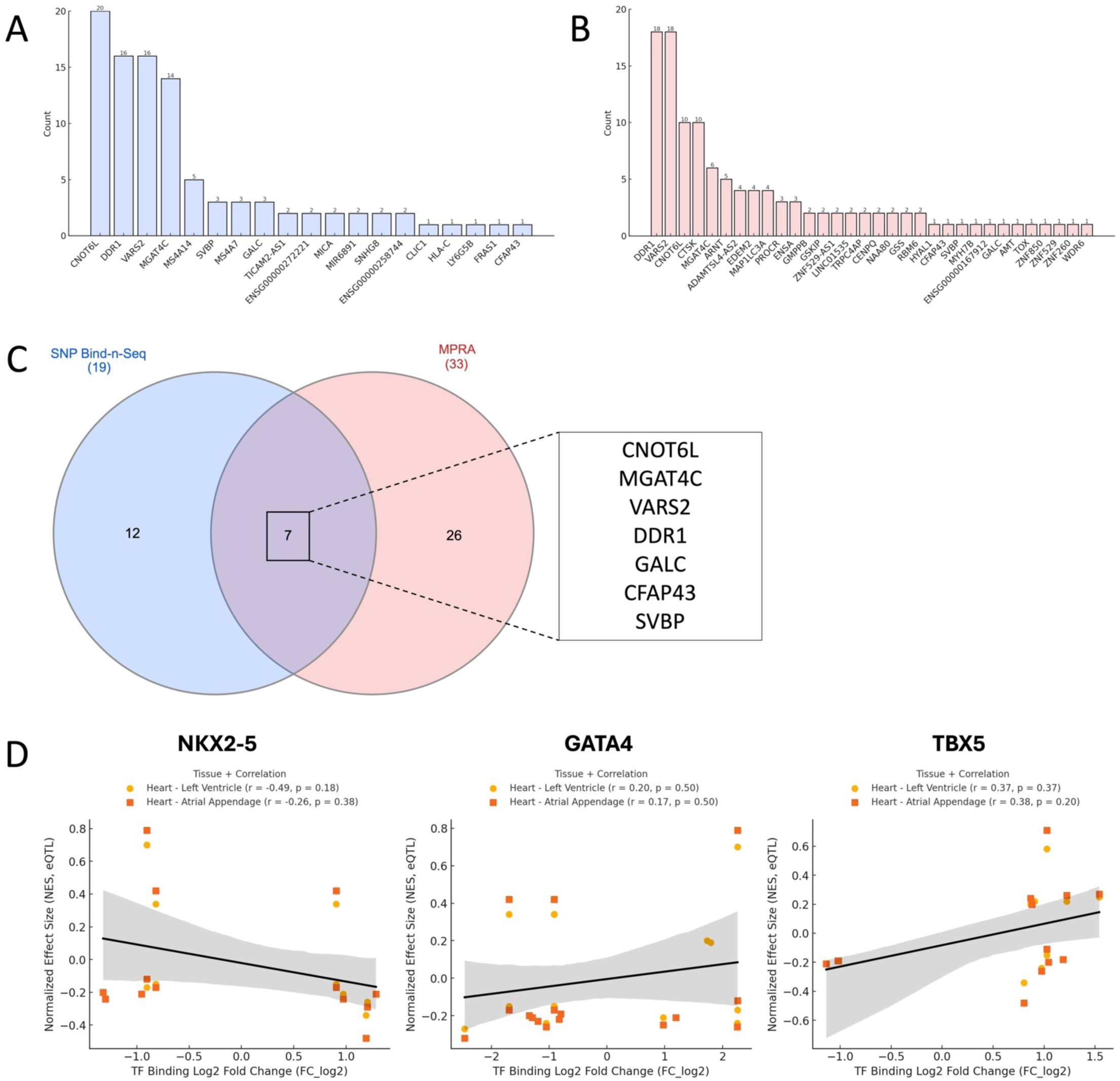
Cardiac eQTL analysis of variants with genotype-dependent regulatory activity in SNP Bind-n-Seq and MPRA. A-B) List of genes (x-axis) and count of variants (y-axis) with cardiac eQTLS from the A) SNP Bind-n-Seq and B) MPRA experiments. C) Venn diagram of genes in cardiac eQTL genes from both experiments. D) Correlation analysis of TF binding fold change and eQTL normalized effect size.

**Supplementary Figure 15:**
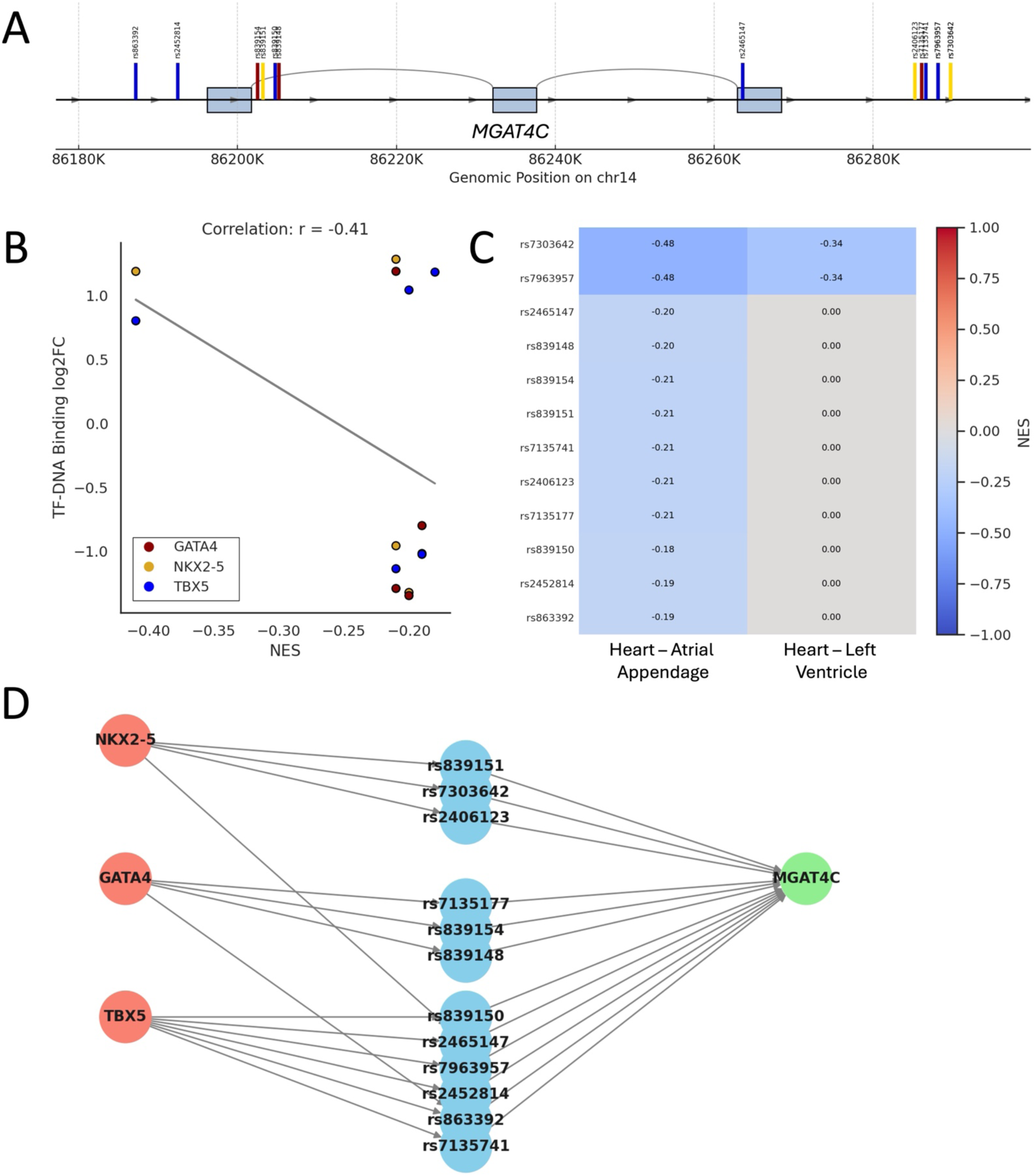
Cardiac eQTL analysis of MGAT4C. **A)** Genomic map of MGAT4C with variants in cardiac eQTL. Variants are displayed as colored lines if they altered NKX2-5 (yellow), GATA4 (red), and TBX5 (blue) binding. **B)** Correlation analysis of cardiac eQTL NES and TF binding fold change. **C)** Heatmap of NES of each variant per tissue. **D**) Interaction network of cardiac eQTL genes, variants, TF with altered binding.

**Supplementary Figure 16:**
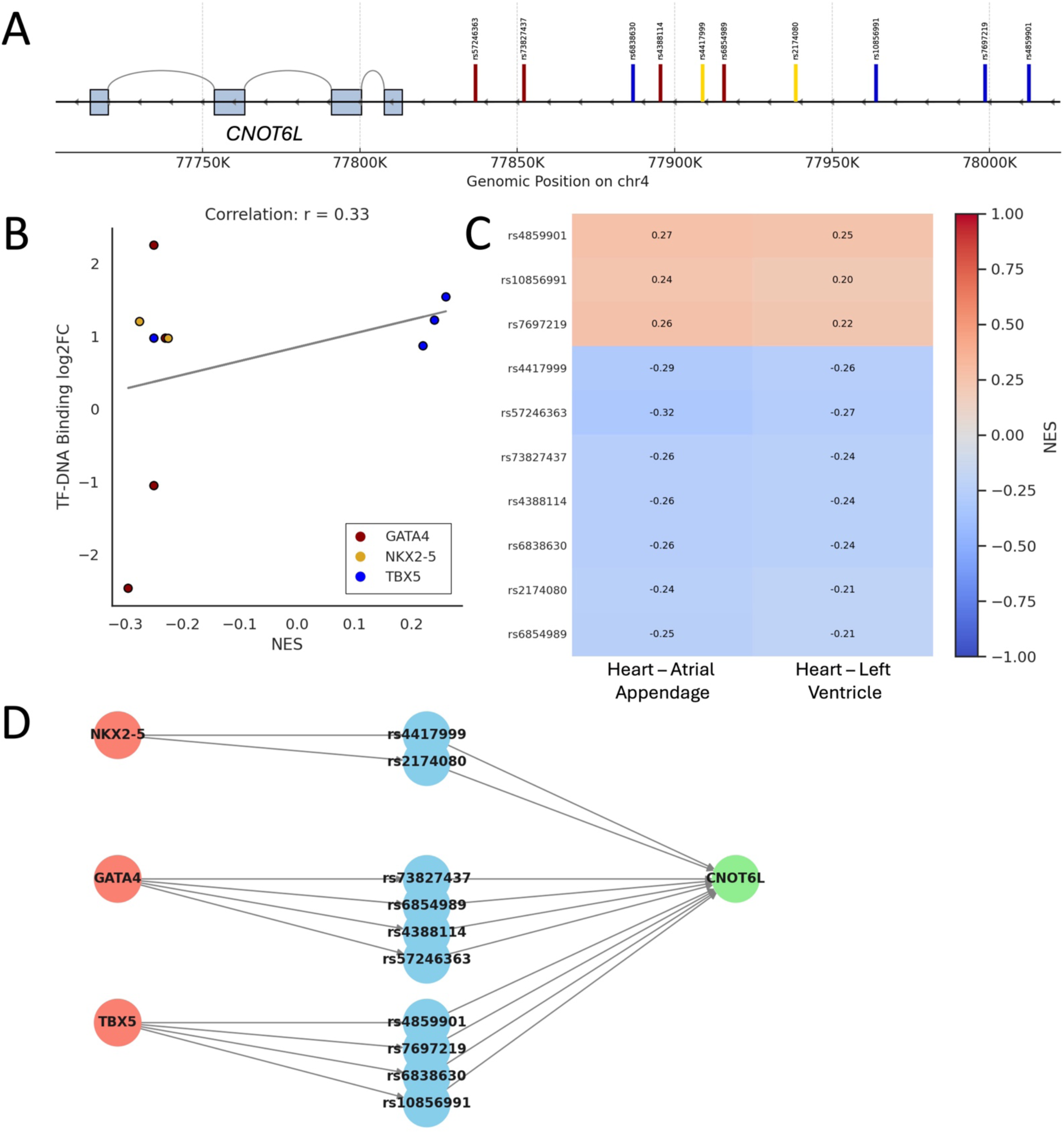
Cardiac eQTL analysis of CNOT6L. **A)** Genomic map of MGAT4C with variants in cardiac eQTL. Variants are displayed as colored lines if they altered NKX2-5 (yellow), GATA4 (red), and TBX5 (blue) binding. **B)** Correlation analysis of cardiac eQTL NES and TF binding fold change. **C)** Heatmap of NES of each variant per tissue. **D**) Interaction network of cardiac eQTL genes, variants, TF with altered binding.

**Supplementary Figure 17:**
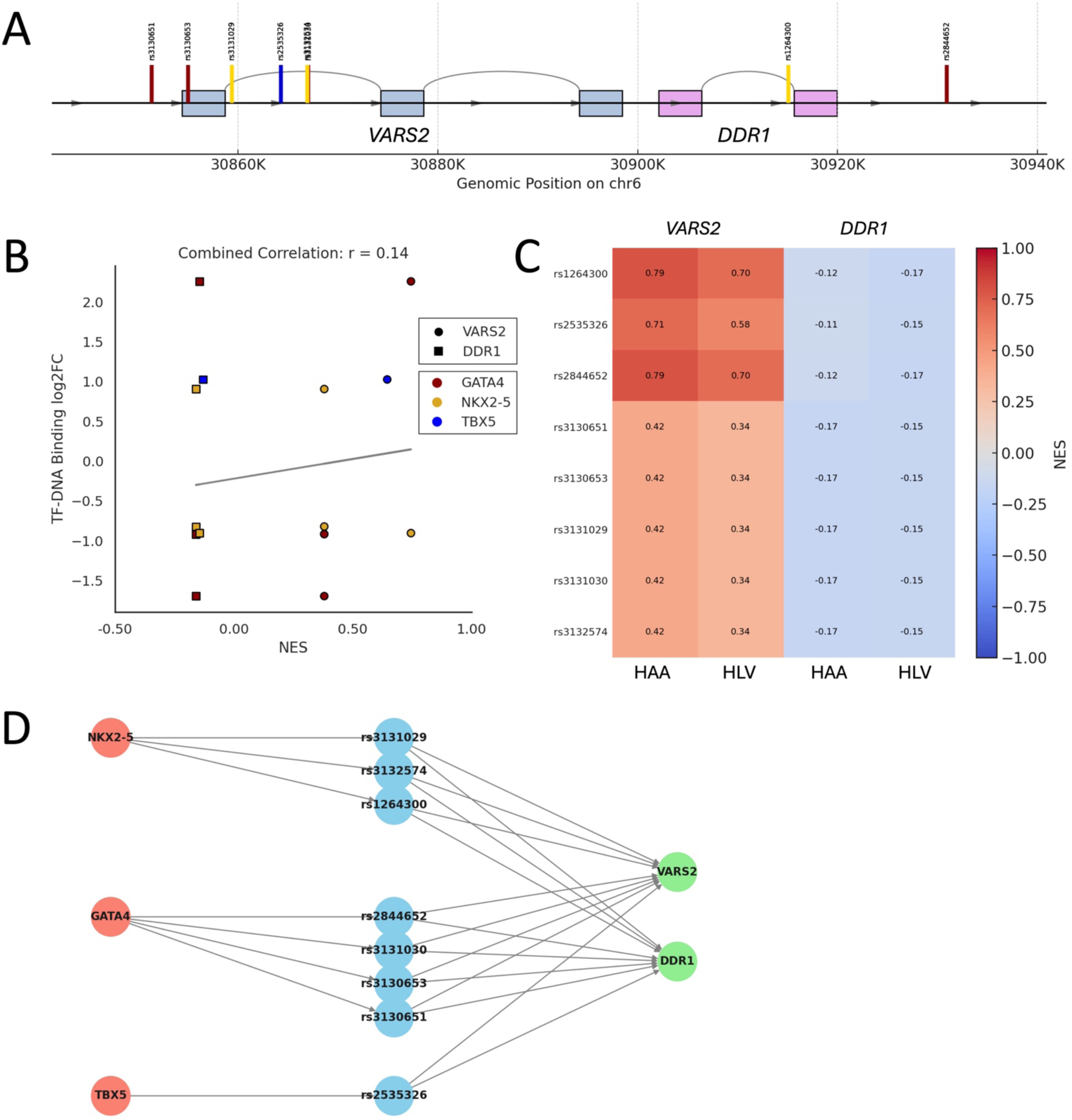
Cardiac eQTL analysis of VARS2 and DDR1. **A)** Genomic map of MGAT4C with variants in cardiac eQTL. Variants are displayed as colored lines if they altered NKX2-5 (yellow), GATA4 (red), and TBX5 (blue) binding. **B)** Correlation analysis of cardiac eQTL NES and TF binding fold change. **C)** Heatmap of NES of each variant per tissue. **D**) Interaction network of cardiac eQTL genes, variants, TF with altered binding.

